# Cardiac-targeted rAAV5-S100A1 gene therapy protects against adverse remodeling and contractile dysfunction in post-ischemic hearts

**DOI:** 10.1101/2023.03.03.529004

**Authors:** Dorothea Kehr, Janek Salatzki, Birgit Krautz, Karl Varadi, Jennifer Birkenstock, Philipp Schlegel, Erhe Gao, Walter J. Koch, Johannes Riffel, Florian André, Karsten Peppel, Hugo Katus, Norbert Frey, Martin Busch, Helga Pfannkuche, Julia Ritterhoff, Andreas Jungmann, Patrick Most

**Affiliations:** Division of Molecular and Translational Cardiology, University Hospital Heidelberg, 69120 Heidelberg, Germany; Department of Internal Medicine III, University Hospital Heidelberg, 69120 Heidelberg, Germany; Center for Translational Medicine, Temple University, Philadelphia, PA 19140, USA; Deutsches Zentrum für Herz- und Kreislaufforschung (GCCR), Partner site Heidelberg/Mannheim; Deutsches Zentrum für Herz- und Kreislaufforschung (GCCR), Partner site Kiel; Department of Cardiology, Robert-Bosch-Hospital, 70376 Stuttgart, Germany; CSL LCC, King of Prussia, Philadelphia, PA 19406, USA; Institute of Veterinary Physiology, Faculty of Veterinary Medicine, Leipzig University, 04104 Leipzig, Germany; Center for Translational Medicine, Jefferson University, Philadelphia, PA 19107, USA

**Author notes:** Address for correspondence: Prof. Dr. Patrick Most, Division of Molecular and Translational Cardiology, Department of Internal Medicine III, Heidelberg University Hospital, INF410, 69120 Heidelberg, Germany.

**Keywords:** Cardiac gene therapy, S100A1, adeno-associated vector, inflammation, heart failure

## Abstract

Toxicity by recombinant adeno-associated viruses (rAAV) in clinical gene therapy trials (e.g., by rAAV9-mediated fatal liver failure) significantly impairs translation of preclinical rAAV-based cardiac gene therapies employing these vectors. For rAAV5 - a capsid that has shown long-term safety in clinical trials - our translational study demonstrates effective transduction of the left ventricle (LV) of healthy pigs via catheter-based retrograde intravenous delivery (CRID) by means of luciferase reporter gene biodistribution analyses. Combination of rAAV5 with the cardioprotective human gene *S100A1* (*hS100A1*) prevents LV myocardial infarct (MI) enlargement and improves LV systolic contractile performance in a porcine model of post-MI chronic cardiac dysfunction. Use of a cardiac-biased promoter ensured the cardiac-directed expression of the therapeutic human transgene without signs of clinical toxicity. The beneficial effects of rAAV5-*hS100A1* were linked to an attenuated activity of post-MI inflammatory gene networks and this was further validated in a murine model. These novel data together with proven scalable producibility and low pre-existing immunity against rAAV5 in humans may collectively advance clinical translation of rAAV5-*hS100A1* as a gene therapy medicinal product (GTMP) for a common cardiovascular disease, such as chronic heart failure (CHF).

**Highlights:**

- Recent fatal adverse events in recombinant adeno-associated virus (AAV)-based clinical gene therapy trials advise the use of rAAV serotypes with proven long-term clinical safety, such as rAAV5, for the pre-clinical development and clinical translation of rAAV-based cardiac gene therapy medicinal products.
- In a biodistribution and therapeutic proof-of-concept study in farm pigs, rAAV5 was identified as an effective viral vector for cardiac gene transfer and gene therapy for post-ischemic cardiac dysfunction when applied by a standardized cardiac-targeted catheter-based route of administration with the luciferase reporter and cardioprotective human gene S100A1 (*hS100A1*), respectively.
- A systems biology analysis linked the novel finding of mitigated inflammatory and activated cardioprotective gene network activities in rAAV5-*hS100A1* treated postischemic myocardium with improved study left ventricular ejection fraction and prevention of myocardial infarct extension, respectively, which warrants further mechanistic molecular studies.
- Since rAAV5 has been recently approved for clinical use in a non-cardiac indication and cardiac-targeted S100A1 gene therapy has been effective in numerous pre-clinical animal models of acute and chronic cardiac dysfunction, our translational data support an expedited developmental path for rAAV5-*hS100A1* throughout investigational new drug-enabling studies towards a first-in-human clinical trial for post-myocardial infarction heart failure.

## Introduction

As a recent technology, gene therapy bears the potential for curative treatments against acquired and hereditary cardiac disorders that remain with unmet clinical needs.^1–3^ But in cardiology the bench-to-bedside iteration has not made the desired clinical impact yet.^4–7^ Instead, other disciplines, such as ophthalmology, neurology, hematology or endocrinology, delivered the first recombinant adeno-associated virus (rAAV)-based gene therapy medicinal products (GTMPs) with European Medical Agency (EMA) marketing authorization. They are directed against hereditary retinal dystrophy (rAAV2-*hRPE65vs*)^8^, spinal muscular atrophy (rAAV9-*SMN1*)^9^, aromatic L-amino acid decarboxylase deficiency (rAAV2-*ddc*)^10^, hemophilia A (rAAV5-*factor VIII*)^11^ or, most recently, hemophilia B (rAAV5-*factor IX).* Hence, conclusions drawn from these clinical programs should advice the selection of appropriate AAV vector types for cardiac GTMP programs to mitigate the risk of failure throughout pre-clinical and clinical development.^12–14^ Of note, rAAV9, which engendered considerable interest as a vector for treating hereditary and common cardiac disorders^1,3^, provoked numerous serious (SAEs) and fatal adverse events (FAEs) in clinical studies for spinal muscular atrophy or Duchenne muscular dystrophy (rAAV9-*miniDMD*)^15,16^. To this rAAV5 contrasts with proven long-term clinical safety and efficacy in patients, such as for hemophilia A or B. But the suitability of rAAV5 for cardiac gene delivery and therapy in human-relevant large animal models with a clinically applicable route of administration (RAO) has not been determined yet.

Given that catheter-based cardiac-targeted gene delivery is currently employed as state-of-the-art route of administration (ROA) in early clinical trials both for an rAAV1-(NCT04179643) and rAAV2-based (NCT04703842) GTMP against congestive heart failure (CHF), we first determined whether rAAV5 is a suitable vector for effective cardiac gene delivery to human-sized hearts in healthy farm pigs by administering an rAAV5-luciferase (luc) vector by a standardized percutaneous catheter-based retrograde cardiac-targeted intravenous delivery (CRID) procedure. These results informed a therapeutic proof-of-concept study using cardiac magnetic resonance (CMR) imaging to probe rAAV5 with the validated CHF target S100A1 as a gene addition treatment with a clinically relevant dosage in farm pigs with post-myocardial infarction (MI) induced chronic systolic dysfunction by CRID. S100A1 was chosen as therapeutic payload given its advanced developmental stage as a GTMP candidate for CHF treatment^17–23^ and the in-depth understanding of its molecular mode of action as a beneficial regulator, e.g., of excitation-contraction (ec) coupling, sarcoplasmic reticulum calcium cycling, of mitochondrial energy supply and myofilament function.^24–34^

Key findings of our translational study not only proved a favorable cardiac transduction pattern for rAAV5 in human-sized hearts when delivered by CRID but unveiled its effectiveness as a therapeutic vector in combination with S100A1 against post-MI chronic detrimental effects by improving the CMR surrogate parameters left ventricular ejection fraction (LVEF) and protecting against LV MI mass increase. Interestingly, by a weighted gene co-expression network analysis (WGCNA) of porcine post-MI myocardial RNA-sequencing data, we discovered a novel function of S100A1 acting as an antagonist of detrimental inflammatory and activator of beneficial cardioprotective gene networks. Overall, we expect that our novel translational data for rAAV5*-hS100A1* will inform and expedite further investigational new drug (IND)-enabling studies towards a first-in-human (FiH) clinical trial. rAAV5s long-term clinical safety profile and excellent scalable manufacturing capabilities may render the vector an excellent candidate for the treatment of a common cardiac disease, such as CHF.

## Methods and Materials

Additional resources are available from the corresponding author upon reasonable request. A more detailed description of each method is provided in the appendix extended methods section. All animal procedures and experiments were carried out according to the ‘Guide for the Care and Use of Laboratory Animals’ (National Institutes of Health) and were approved by the local Institutional Animal Care and Use Committee of Baden-Württemberg, Germany and Jefferson University, PH, USA. All mice were housed at 22°C with a 12-hour light, 12-hour dark cycle with free access to water and standard chow. Bay cage types are used for housing of pigs. Per pig, a body weight prescribed floor space of at least 0.5 and 0.7 m^2^ were provided according to directive 2010/63/EU with straw bedding. The environment was enriched by pellet balls, chains and gnawing rods. Humidity and temperature were kept at 50-60 % and 20-24 °C, respectively. Access to water was unlimited and restricted food provided twice/d (SAF130M).

### rAAV manufacturing

rAAV vectors were produced, purified, titrated, and stored as described previously.^17,20,35^

### Cardiac-targeted gene delivery to porcine hearts

Cardiac-targeted catheter-based retrograde intravenous delivery (CRID) of rAAV vectors both to normal and infarcted pig hearts was carried out as described elsewhere.^17,20,35^

### Tissue vector abundance, gene expression and immunoblot analyses

Isolation of tissue DNA and RNA including reverse transcription and PCR-based vector genome abundance and gene expression quantification as well as immunoblotting was conducted as described previously.^17,20,35^

### Pig myocardial infarction model

Post-myocardial infarction cardiac dysfunction in male German farm pigs was induced by percutaneous transluminal temporary occlusion of the left circumflex artery (LCX) as described elsewhere.^17,20,35^

### Cardiac magnetic resonance and imaging analysis

Assessment of left ventricular (LV) ejection fraction (EF) and myocardial infarct (MI) size was performed by published standard cardiac magnetic resonance protocols adopted to porcine anatomy.^36–38^

### RNA-sequencing of porcine myocardium and weighted gene correlation network analysis

Next-generation sequencing of isolated myocardial LV RNA and subsequent weighted gene correlation network analysis (WGCNA) was carried out as described elsewhere.^39–41^

### Mouse myocardial infarction model and intramyocardial injections

Induction of myocardial infarction by experimental ligation of the left anterior descending (LAD) artery and intramyocardial injections of rAAV5 vectors were performed as described previously.^34,39,42^

### Mouse echocardiography

Transthoracic two-dimensional echocardiography of post-infarction murine hearts was conducted as described elsewhere.^34,42^

### Statistics

The numbers of independent experiments/ animals are specified in the relevant figure legends. Data are expressed as mean + standard error of the mean (SEM). Statistical analysis was performed with Prism 8.0 or 9.0 software (GraphPad). Normal distribution of data was verified by Shapiro-Wilk test. For normal distributed data, statistical comparisons between 2 groups were conducted by unpaired, two-tailed t-test. Statistical comparisons between 3 or more groups were conducted by one-way or two-way ANOVA followed by a Tukey posthoc analysis to determine statistical significance. For non-normal distributed data Kruskal-Wallis test was performed between 3 or more groups with subsequent Dunn’s test for multiple testing. For comparison over time, Bonferroni posthoc analysis was performed. The value of p < 0.05 was considered statistically significant.

## Results

### Myocardial distribution of rAAV5, rAAV6 and rAAV9 after cardiac-targeted retrograde intravenous administration in pigs

We first investigated whether cardiac-targeted catheter-based retrograde intravenous delivery (CRID) of rAAV5 - as a clinically applicable ROA - might result in a suitable *in vivo* cardiac transduction pattern of the porcine left heart. These results were systematically compared against rAAV6 and rAAV9 in our biodistribution study (Figure 1A) given their previously demonstrated ability to transduce diseased porcine hearts by CRID, e.g., with *hS100A1* and other transgenes.^17,20,35, 43–45^ Each vector was given with a dosage of 1×10^13^ vector genome copies (vgcs) carried the *luciferase* (*luc*) reporter gene under control of a synthetic cardiac-biased expression element utilizing a myosin light chain ventricle-2 (MLCv2) promoter fragment as described previously.^22,25,40^ To achieve a standardized CRID procedure (Figure 1B), we implemented quality measures for the experimenter further detailed in the extended methods section and shown by appendix figure A1. Figure 1C to 1F depict the key findings for the cardiac and extra-cardiac biodistribution of rAAV5, rAAV6 and rAAV9 by systematic luminometric assessment of tissue *luc* activity after 30 days. rAAV5 showed a significantly higher reporter activity than rAAV9 in each of the three myocardial segments along the apical-basal axis of the porcine left ventricle (LV) (Figure 1C). In pooled samples of the three myocardial segments (apical, medial and basal) per animal, rAAV5-treated hearts yielded an approximately 30-fold greater reporter activity than rAAV9 (rAAV5 167.179±16.083 RLU/mg tissue protein vs. rAAV9 5.084±835 RLU/mg tissue protein; data are given as mean±SEM, P<0.01 rAAV5 vs. rAAV9, n=5 pooled myocardial samples per group). rAAV6 showed the highest but most heterogenous myocardial *luc* activity pattern of the tested vectors with an almost 10-fold difference between the heart’s apex and basis (Fig. 1D). Subsequent extra-cardiac measurements of *luc* reporter activity demonstrated that rAAV6 yielded the highest absolute off-target reporter activities in all organs (Fig. 1E), while rAAV9 yielded the greatest relative off-target reporter activity when normalized to its cardiac *luc* activity levels (Fig. 1F). From these data, we concluded that rAAV5 is an effective vector for cardiac-targeted gene delivery via our CRID SOP in a large animal model that closely approximates human cardiac anatomy. Subsequent CRID of 1×10^13^ vgcs of rAAV5 - carrying the *hS100A1* gene under control of the MLCv2 fragment as described previously^22,25^ - to porcine hearts resulted in a robust myocardial S100A1 protein expression increase after 30 days compared to rAAV5-*luc* control (n=4 animals per group) (appendix figure A3). These results subsequently informed the design of an rAAV5-based gene therapy study in a post-myocardial infarction (post-MI) pig model to determine its suitability as a therapeutic vector in a clinically relevant setting using the *hS100A1* transgene.

**Fig. 1A.**
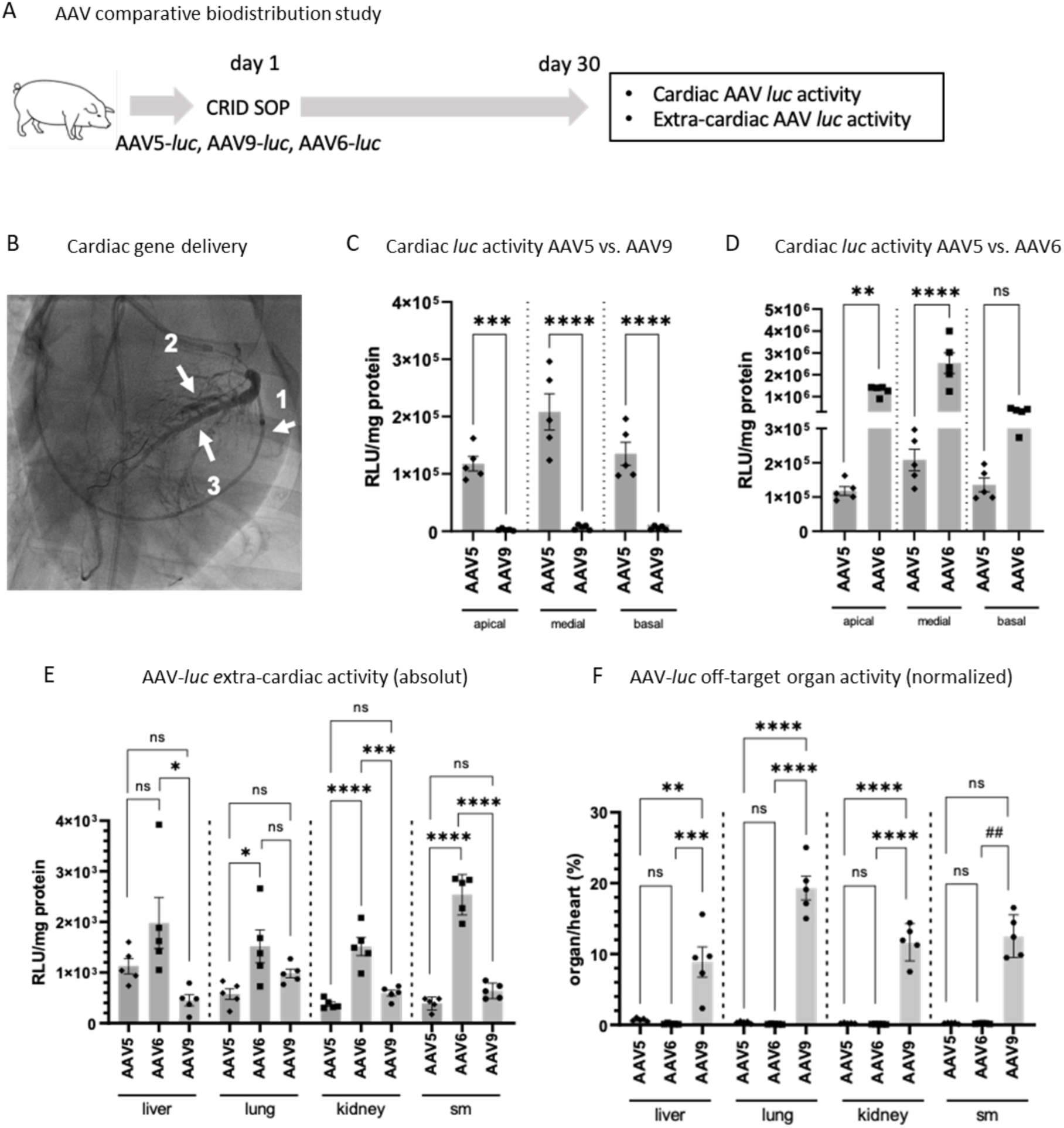
shows the comparative 30-day biodistribution study design. **Fig. 1B** depicts a representative fluoroscopic image as part of the standardized CRID SOP. Arrow 1: appropriate position of the retroinfusion catheter in the coronary sinus with subsequent infusion into the cardiac anterior interventricular vein (AIV); arrow 2: balloon catheter in the left anterior descending artery (balloon inflated at the time of the image); arrow 3: retrograde installation of contrast agent in the AIV via the retroperfusion catheter to ascertain the correct position and sufficient blocking of the retroperfusion catheter tip prior to CRID. **Fig. 1C and D** illustrate the *luc* reporter gene activity along the apico-basal axis of the healthy porcine heart 30 days after CRID of rAAV5-*luc* compared with rAAV6-*luc* and rAAV9-*luc*, respectively. **Fig. 1E and F** display the absolute extrac-cardiac and normalized off-target organ gene activity levels for each rAAV serotype, respectively, 30 days after the CRID SOP. *Luc* reporter gene activity is given as relative light units (RLU) in relation to mg of extracted tissue protein. ****P<0.001; ***P<0.01; **P<0.03, *P<0.05. Data are presented as mean+-SEM. n=5 animals per group. Statistical comparisons were conducted by one-way ANOVA, followed by Sídák‘s multiple comparison test (C, D), followed by Tukey‘s posthoc analysis (E, F) and conducted by Kruskal-Wallis test with Dunn‘s multiple comparisons test for F (skeletal muscle).

### Impact of rAAV5-*hS100A1* on cardiac performance and damage in the cardiac injury pig model assessed by CMR

Following published expert consensus opinion, we first implemented cardiac magnetic resonance (CMR) as non-invasive gold standard for the assessment of left ventricular (LV) ejection fraction (EF) and LVMI changes in the employed post-MI systolic dysfunction porcine model,^17,20,35^ as recommended CMR surrogate parameters with hard clinical outcome.^46–48^ Standardized CMR acquisition sequences were adopted to the porcine cardiac anatomy and utilized for the serial measurement of LVEF and LVMI size.^36–38^ and the CMR analysis was blinded to the treatment type. LVMI size was quantified by late gadolinium enhancement (LGE) and, additionally assessed by global native T1-relaxation 2 weeks after experimental MI before study enrollment. Only animals with an LVMI size >14% resulted in a significant decline in LVEF (appendix figure B1) and were therefore assigned to the rAAV5-*luc* control and rAAV5-*hS100A1* groups of the intervention study (Figure 2A) to receive a single dosage of 1×10^13^ vgcs of the respective vectors by the CRID SOP as previously described.^22,25^ Prior to rAAV5-based gene therapy, the average LVMI size did not differ significantly between both groups (appendix figure B2). rAAV5-*hS100A1* post-MI treatment prevented the extension of total LVMI mass that had occurred in all animals of the control group assessed at end of the 3-month follow-up period (appendix figure B3) both by LGE signal and T1 relaxation time (Figure 2B and C). Concurrent CMR-based analysis of LVEF changes unveiled a significant increase of LVEF in the rAAV5-*hS100A1* treated arm over the control group (Figure 2D), which is in line with previous results from studies using viral-based *hS100A1* gene therapy in small and large animal post-MI heart failure models.^22–25,27–28,73,80^ Heart rate, LV end-diastolic and end-systolic volumes, respectively, are given in appendix figure B4. From this outcome, we concluded that rAAV5 is a suitable rAAV serotype to sufficiently deliver the *hS100A1* gene to human-sized diseased hearts and subsequently entail long-term improvement of cardiac performance. But the yet unreported protective action of rAAV5-*hS100A1* against chronic LVMI extension prompted additional molecular studies.

**Fig. 2A.**
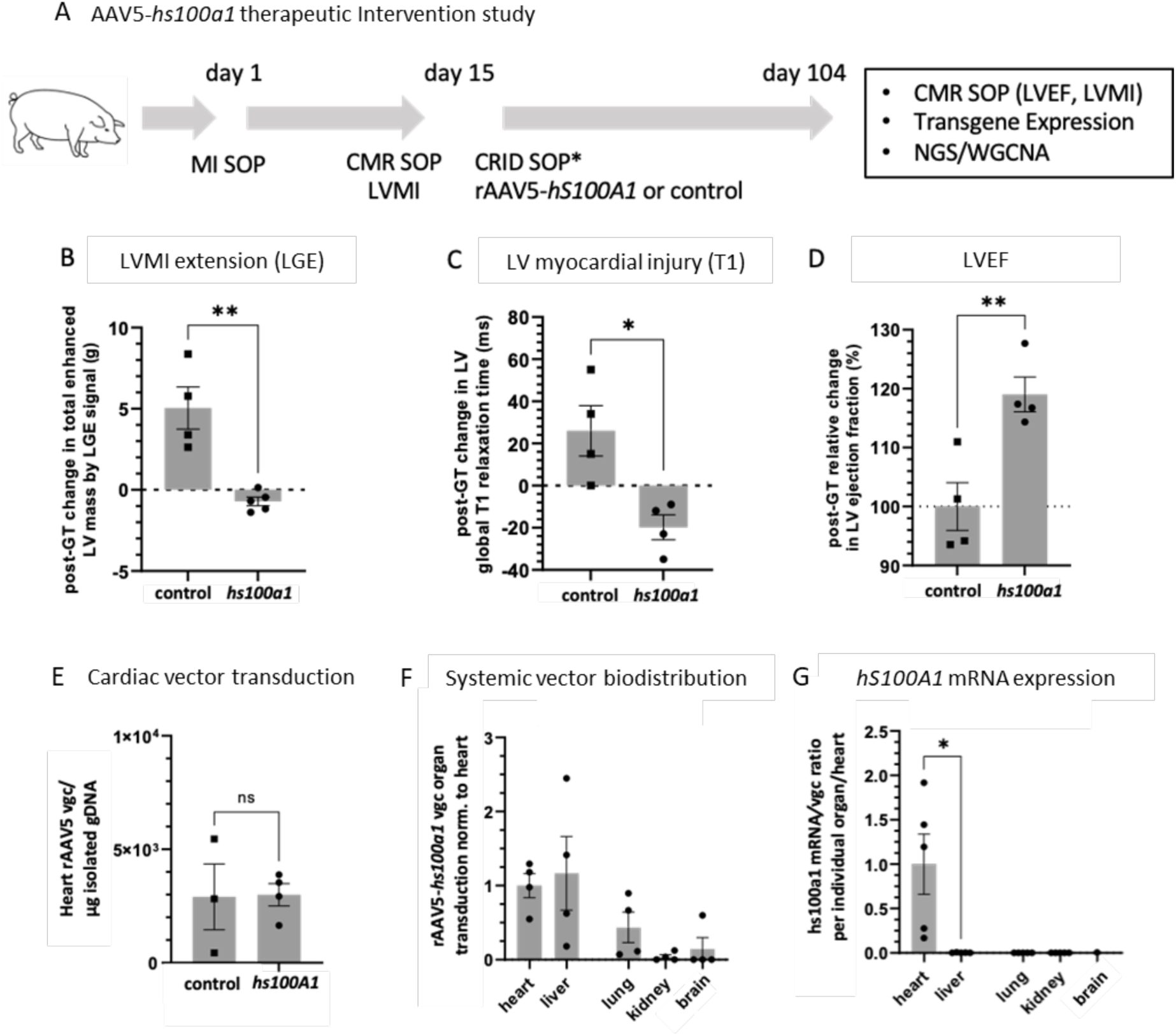
describes the design of the intervention study assessing the therapeutic impact of rAAV5*-hS100A1* in the pig post-MI cardiac dysfunction model. **Fig. 2B and 2C** depict the rAAV5-*hS100A1*-mediated post-gene therapy (post-GT) protection against chronic LVMI mass extension compared with the control group assessed both by LGE signal (g) and global LV T1 relaxation time (ms). Data are given as individual post-GT change for each animal (day 15 to 104). LGE **P<0.03; control n=4 and *hS100A1* n=5; T1 *P<0.05; control n=4 and *hS100A1* n=4. **Fig. 2D** depicts the significant post-GT improvement of LVEF in the AAV5-*hS100A1* versus the control treated group. Data are given as relative LVEF 3-month post-GT normalized to control. **P<0.03; n=4 and *hs100a1* n=4. **Fig. 2E** shows equal transduction of the porcine heart by the control rAAV5 (n=3) and rAAV5-*hS100A1* vector (n=4) 3-months post-CRID SOP. Data are shown as vector genome copy (vgc) numbers per μg of isolated genomic DNA (gDNA) from the porcine heart for each group. **Fig. 2F** displays the systemic distribution of the rAAV5-*hS100A1* vector (n=4) 3-months post-CRID in a representative set of parenchymal organs. Data are presented as individual organ/heart ratio of the respective organ rAAV5*-hS100A1* vector genome copy (vgc) number/μg genomic DNA. **Fig. 2G** highlights the cardiac expression of *hs100a1* mRNA versus liver and other organs. *P<0.05. Data are given as ratio of *hS100A1* mRNA copy numbers normalized to the heart‘s per rAAV5-*hS100A1* vgc value. *hS100A1* mRNA tissue copy numbers were calibrated using an *hS100A1* gene-containing plasmid standard curve. Data are presented as mean+-SEM. Statistical comparisons were conducted by unpaired two-tailed t-test (B, C, D, G) and by Mann Whitney test (E).

### S100A1 expression analyses, exploratory safety, and toxicity assessment after AAV5-*hS100A1* cardiac-targeted delivery

To confirm an effective myocardial gene delivery by cardiac-targeted CRID and characterize the biodistribution of rAAV5 and cardiac expression of the therapeutic *hS100A1* transgene, we assessed vgcs in different organs and distinguished between human and porcine *S100A1* gene expression as described previously.^17,20,35^ We first confirmed that rAAV5 vgc numbers in control and rAAV5-*hS100A1* treated hearts are on comparable levels (Figure 2E). Also, we performed analysis of extra-cardiac *hS100A1* expression analysis detecting rAAV5-*hS100A1* genomic DNA, e.g., in liver, lung, kidney or brain (Figure 5C) indicating that the cardiac-biased promoter ensured predominant *hS100A1* mRNA expression in rAAV5-*hS100A1* treated hearts (Figure 2F). Regular inspections of animal behavior, body temperature as well as food intake and excrement did not yield any abnormalities within the 3-month post-treatment observation period (data not shown). White and red blood cell parameter as well as platelet count and coagulation parameters were just as well within porcine laboratory standard ranges. Likewise, electrolytes and blood glucose levels, and particularly biomarkers for liver integrity and synthesis as well as kidney function and pancreas integrity did not show any abnormalities (appendix C1). Finally, the cardiotoxicity protocol involving troponin T (appendix figure C1) and ECG parameters (data not shown) did not deliver any sign of cardiac damage or disturbed repolarization. From these results, we concluded that rAAV5 together with catheter-based CRID is suitable to deliver sufficient *hS100A1* gene copies to the diseased pig heart to exert the demonstrated therapeutic effect on LVEF and chronic MI extension.

### Weighted gene co-expression network analysis indicates active cardioprotective and attenuation of inflammatory gene programs after AAV5-*hS100A1* based gene therapy

Aiming at underlying molecular clues for the rAAV5-*hS100A1* post-GT superior study outcome, we next carried out bulk RNA-sequencing 3-months after gene transfer from LV myocardial tissue samples. Then, a weighted gene correlation network analysis (WGCNA) was performed by applying a recently published WGCNA R software package to capture clusters of highly co-expressed genes (modules) with LVMI and LVEF changes.^41^ Only module eigengenes (MEs) with a significant and strong correlation (>0.8 and <-0.8) with LVMI and LVEF changes were further examined. The principal component (PC) analysis prior to WGCNA segregated rAAV5-*luc* from the rAAV5-*hS100A1*-treated animals (appendix figure D). Figure 3A illustrates the module-trait relationship matrix depicting a significant and strong negative correlation (>0.8 and <-0.8) of the turquoise and cyan MEs with LVEF and chronic LVMI extension, respectively. These were further processed by *in-silico* pathway databases (Figure 3A), such as Reactome, GO and KEGG, to infer molecular pathways with potential mechanistic relevance for rAAV5-*hS100A1* mediated therapeutic effects. As such, our *in-silico* examination yielded a significant overrepresentation of the inflammatory and immunological signaling pathways in the turquoise ME versus the LVEF state inferring a lower activity of neutrophil as well as T- and B-cell related gene networks, such as “*neutrophil degranulation”*, “*T cell receptor signaling”*, “*signaling by the B cell receptor”*, “*downstream signaling events of B cell receptor”, “Fc epsilon receptor signaling”* and *“Fc epsilon receptor signaling mediated NF-kB activation”* (Figure 3B) in rAAV5-*hS100A1* treated porcine hearts where post-MI LVEF was significantly enhanced. In addition, a lower activity of *“S-phase”*, *“Synthesis of DNA”* and *“Mitotic G1 phase and G1/S transition”*, representing the cellular commitment to DNA synthesis, may reflect in part the previously reported anti-hypertrophic effect of S100A1 in diseased myocardium.^23,25,26,28,39,80^ Further functional pathway annotation unveiled a significant overrepresentation of the pathways “*calcium signaling”*, “*mitochondrial translation initiation, elongation and termination”* as well as “*ATF4 activates genes in response to endoplasmic reticulum stress”*, “*PERK regulates gene expression”* and “*unfolded protein response”* as well as “*signaling by retinoic acid”* and “*fibroblast growth factor receptor 2 ligand binding and activation”* pathways in the cyan ME that negatively correlates with the changes in the chronic LVMI extension CMR surrogate global T1 (Figure 3C and D) and, in turn, indicates a higher activity of these cardioprotective pathways in rAAV5-*hS100A1* treated myocardium. From this we concluded that the therapeutic actions of S100A1 in dysfunctional myocardium may exceed the previously described direct regulation of calcium cycling, myofilament and energy homeostasis effector genes by us and others^49,50^ and potentially extend to the modulation of cardioprotective transcriptional programs by a yet unknown molecular mechanism.

**Fig. 3A.**
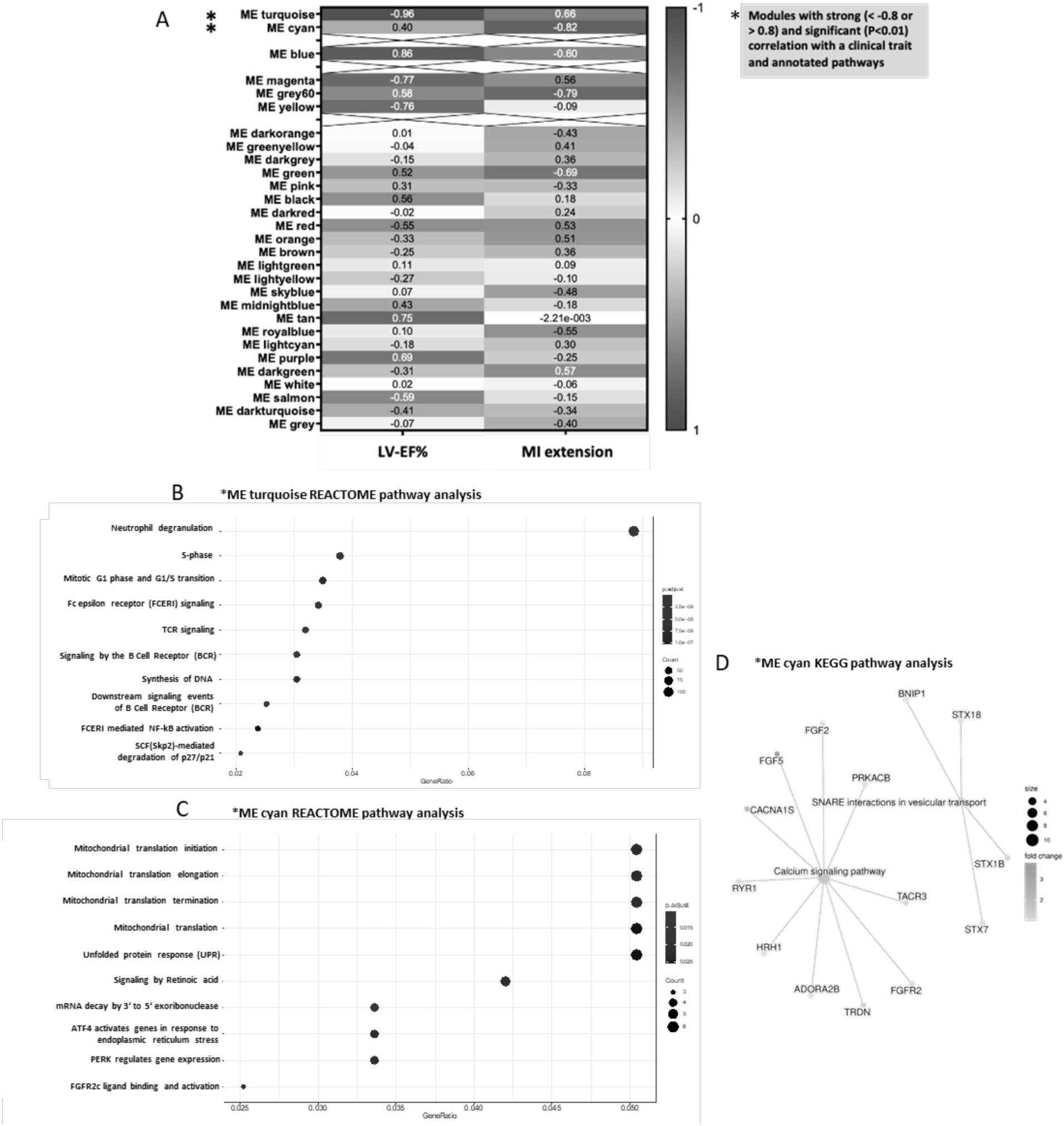
shows the result of the WGCNA for the two clinically relevant surrogate parameters LVEF and chronic LVMI extension, where ME turquoise and ME cyan strongly correlate with LVEF changes (P<0.001) and LVMI extension (P<0.03), respectively. **Fig. 3B** highlights the annotated and significantly enriched pathways in the ME turquoise that negatively correlates with the difference in LVEF. The mostly proinflammatory gene ensembles - that negatively correlate with LVEF reflect activation of the innate and adaptive immune system in the post-MI pig heart - display a lower expression in the rAAV5-*hS100A1* group compared with control (rAAV5-*hS100A1 vs.* control n-fold gene expression ratios were < 1). **Fig 3C** and **D** show the annotated and significantly enriched pathways in the ME cyan that negatively correlates with LVMI extension. The mostly energy performance, cardioprotective and calcium signaling pathway gene ensembles - that negatively correlate with LVMI extension - display a higher expression in the rAAV5-*hS100A1* group compared with control since the rAAV5-*hS100A1* vs. control n-fold gene expression ratio were > 1.

### rAAV5-*hS100A1* treatment of LV infarcted mice recapitulates attenuated innate and adaptive immune system activity in myocardium

We subsequently employed a previously published experimental murine MI model^34,42^ with the aim to validate transcriptional key findings inferred from the inflammatory signaling pathway analysis in our large animal post-MI model (Figure 4A). To this end, a total dosage of 2×10^11^ vgcs of either rAAV5-*hS100A1* or AAV5-green fluorescent protein (*gfp)* was injected into the LV wall of the extra-thoraxically exposed heart (Figure 4B) prior the ligation of the LAD. The impact on post-MI LV performance, MI size and gene expression in remote LV myocardium was determined 4 weeks after treatment as previously described.^34,42,51^ Since an effect of S100A1 gene therapy on the immune system activity in remodeled and dysfunctional hearts had not yet been described, we focused on marker genes related to innate and adaptive immune system activity. Figure 4D shows that rAAV5-*hS100A1* treatment improved LV fractional shortening (FS) in the murine model compared to the control groups after 4 weeks. Immunoblot analysis of S100A1 and GFP and protein levels yielded robust S100A1 overexpression in rAAV5-*hS100A1* treated mice with equal GFP levels in both groups (appendix E). The estimated MI size in the rAAV5-*hS100A1* treated hearts was significantly smaller than in the control group (13.48±0.5% vs. 22.63±1.2% scar/LV myocardium; data are given as mean±SEM, rAAV5-*hS100A1* vs. rAAV5-*gfp* P<0.01; n=7 animals per group). Of note, post-MI mice with rAAV5-*hS100A1* treatment exhibited lower levels of various marker genes for myocardial macrophage, neutrophil and T-cell presence and proinflammatory cytokines, including, e.g., *cd68, cxcr2* and *cd4* as well as *il-1b, tnf* and *inf-g* compared to control (Figure 4D). Clearly, these transcriptional findings prompt further mechanistic examinations beyond our first validation in a small animal model given the emerging relevance, e.g., of inflammatory processes in the failing heart that may be beneficially targetable by rAAV5-*hS100A1* gene therapy.

**Fig. 4A.**
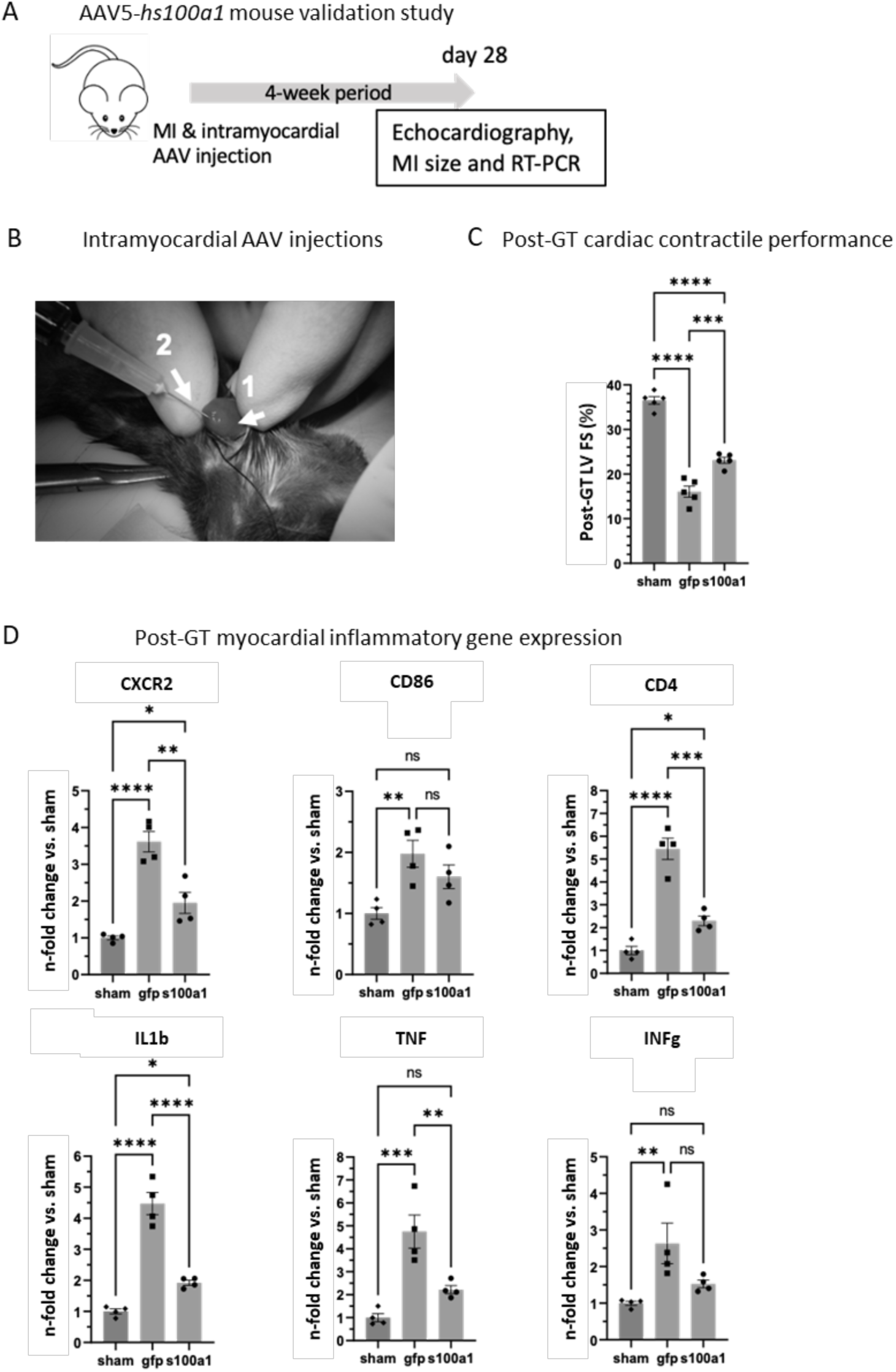
describes the design of the rAAV5-*hS100A1* validation study using intramyocardial injections immediately after experimental MI. **Fig. 4B** shows the published method^39,47^ by which the murine heart was extra-thoraxically exposed (arrow 1) to convey intramyocardial injections (arrow 2) prior to experimental MI procedure by permanent LAD ligation. **Fig. 4C** displays the improvement in LVFS by rAAV5-*hS100A1* compared to control (rAAV5-*gfp*). **Fig. 4D** unveils the attenuation of selected cellular inflammatory gene marker (CXCR2, CD86 and CD4) and mitigation of inflammatory cytokine expression (Il1b, TNF and INFg) in the rAAV5-*hS100A1* treated post-MI myocardium compared to both control (sham and rAAV5*-gfp)* groups. Values were normalized to beta-actin levels. ****P<0.001; ***P<0.01; **P<0.03, *P<0.05; n=4 animals per group. Data are presented as mean+-SEM. Statistical comparisons were conducted by one-way ANOVA, followed by Tukey‘s posthoc analysis (C, D) and by Kruskal-Wallis test with Dunn‘s multiple comparisons test for INFg.

## Discussion

The long-term safety of rAAV5 in clinical trials^54^ renders it an interesting candidate for cardiac GTMP development. In addition, the long-lasting therapeutic efficacy in patients despite preexisting neutralizing antibodies^54^ and scalable producibility^55^ may further predestine rAAV5 as a vector for common cardiovascular indications, such as CHF. This contrasts, e.g., with rAAV9 causing various SAEs and fatal liver failure when employed as a therapeutic vector in various clinical and preclinical studies.^1,3,15,16^ rAAV9’s hepatotoxic potential due to its strong hepatotropism gains further relevance when a therapeutic use of rAAV9 may be considered in critically ill CHF patients. They typically exhibit a commonness of congestive hepatopathy which render the organ even more susceptible, e.g., for rAAV9-mediated toxicity.^56,57^ As such, the most recent FDA approvals of rAAV5-based gene therapies for a hereditary hepatic bleeding disorder strongly supports the notion that this serotype could be a low-risk vector for future human cardiac GTMP development. Hence, we first tested the suitability and efficacy of rAAV5 for cardiac gene delivery to a human-sized heart with a clinically applicable ROA in a porcine model that roots in the percutaneous retrograde coronary venous angiography procedure. To ensure safety and reproducibility, we implemented specific quality measures to standardize our CRID procedure, including prevention of leakage during vector delivery, avoidance of accidental selective infusion into distal coronary venous branches and confirmation of vascular integrity after gene delivery. Based on our experience, such standardized checkpoints prove particularly useful to deliver high-quality data sets for IND-enabling studies, e.g., by clinical research organizations.

Delivered by CRID, rAAV5 almost evenly transduced the porcine heart along the basic-apical axis like rAAV9 but produced significantly higher myocardial reporter gene expression levels. They were lower than rAAV6 but did not exhibit rAAV6s inhomogeneous cardiac expression pattern as found in our study and in previous reports^17,35,58^. Moreover, rAAV5 both displayed the lowest absolute and relative extra-cardiac reporter gene expression values, which again contrasts with rAAV9 that showed the highest off-target reporter gene expression normalized to its cardiac activity despite the use of the same cardiac-biased expression control element in all tested vectors. This is likely caused by the broad organ tropism of rAAV9 (e.g., toward liver, lung, heart or kidney besides other organs) resulting in high off-target organ transduction irrespective of a systemic or cardiac-targeted administration, as reported previously.^59,60^ Clues for the distinct cardiac and off-target expression characteristics of each serotype may first come from the different abundance of common (e.g., the wild type (wt)AAV receptor KIAAA0319L)^61^ and serotype-specific wtAAV receptor constellations (e.g., wtAAV9: laminin receptor and N-linked glycans; wtAAV6: N-linked sialic acid and EGF receptor; wtAAV5: N-linked sialic acid and PDGF receptor) on the various organs that convey cellular AAV binding and internalization.^62,63^ As additional factors such as pre-existing neutralizing antibodies or interacting serum factors^64,65^ can influence the *in vivo* transduction properties of AAVs, a systematic assessment across different mammalian model systems would be needed to provide a conclusive understanding of this complex mechanistic process. We nevertheless conclude up to this point that rAAV5 provides a suitable cardiac transduction pattern with negligible off-target effects when delivered to a human-sized heart by a clinically applicable ROA in combination with cardiac-biased expression control. Considering the previously reported low efficacy of myocardial gene delivery, e.g., by antegrade intracoronary delivery in the CUPID trial series^1,4,5^, CRID may provide a useful alternative when carried out by invasive cardiologists.

These data eventually prepared the groundwork for a translational proof-of-concept study demonstrating that the favorable to cardiac expression characteristics of rAAV5 can be therapeutically utilized in combination with the validated target *hS100A1* in a large animal disease model that closely approximates human cardiovascular pathophysiology. The previously reported therapeutic effect sizes of the *hS100A1* gene either with rAAV9 or rAAV6 in post-ischemic pig and rodent models^17–20,22,66^ allowed us to use the fewest animals possible to show a similar beneficial impact of rAAV5-*hS100A1* on contractile performance and capture the novel finding that rAAV5-*hS100A1* can prevent LV MI mass enlargement as part of adverse cardiac remodeling. Previous experimental and computational studies on this pathology suggest that this chronic enlargement of the infarcted myocardium is likely driven by a non-ischemic pathomechanism,^67–70^ which is in line with the otherwise intact myocardial perfusion in our porcine model. Depressed contractility in the adjacent zone to the infarcted myocardium seems to subject cardiomyocytes to augmented wall stress leading to perpetuated low-level cardiomyocyte apoptosis and replacement by fibrotic tissue with subsequent reduction of adjacent zone contractility.^67^ Apparently, this vicious cycle has the potential to progressively extend the necrotic myocardial tissue by drawing remote, otherwise healthy myocardium into the low-contractility adjacent zone.^67–70^ Since recent studies linked improved border zone contractile performance to an attenuation of MI extension, S100A1’s previously reported enhancement of post-MI contractile performance, energy restoration, protection from cardiomyocyte apoptosis and mitochondrial protection in vivo and in vitro^19–22,71^ aligns well with the herein documented prevention of chronic MI extension by AAV5-*hS100A1* treatment. Our finding gains further clinical relevance as in patients, non-ischemic MI extension seems to occur in up to 9% of individuals after acute MI with a hospital mortality more than four times higher than MI without extension.^72^ But the incidence and long-term risk of this pathology in the chronic phase after MI is still largely unknown and requires further research.

This result subsequently prompted a cardiac transcriptomic-based WGCNA and enrichment analysis to infer gene networks that may play a mechanistic role in rAAV5-*hS100A1’s* beneficial actions. Most recent work by us on the porcine cardiac transcriptome facilitated the annotation of the retrieved myocardial transcripts.^39,40^ The module-trait relationships (MTRs) were subsequently calculated by pairwise correlation of each module’s eigengene (ME) with the post-MI directed changes in LVMI mass extension and LVEF, respectively. By this, a higher expression and activity of gene networks and components in rAAV5-*hS100A1* treated myocardium could be inferred by our WGCNA that uphold myocardial calcium cycling, benefit mitochondrial energy supply and entail cardioprotection. In this regard, triadin upregulation in rAAV5-*hS100A1* treated hearts, which is a critical factor for structural and functional integrity of the cardiomyocyte calcium release unit,^73^ might synergize with the previously reported enhancement of cardiac excitation-contraction(ec)-coupling by direct molecular actions of S100A1 on RyR2 and SERCA2a. In addition, the previously demonstrated improvement of cardiomyocyte energy supply by enhanced activity of mitochondrial complex V and sealing of the mitochondrial permeability transition pore by S100A1^21,25^ might concur with augmented mitochondrial ribosomes and ATF4-mediated mitochondrial stress response^74^ pathway activity after rAAV5-*hS100A1* treatment. Finally, the previously reported anti-apoptotic and cardioprotective effect of S100A1^34^ aligns well with enhanced activity of cardioprotective FGF-2 and -5 as well as retinoic acid gene ensembles in rAAV5-*hS100A1* treated myocardium, which have been linked to the mitigation of MI size, augmented post-MI pro-angiogenic signaling and abrogated myocardial oxidative stress.^75–77^ Clearly, the notion of rAAV5-*hS100A1* activated cardioprotective gene networks, as inferred by our WGCNA, warrants further mechanistic studies but our results nevertheless suggest that the impact of S100A1 on several protective gene pathways could complement the otherwise previously reported beneficial molecular actions of the EF-hand Ca^2+^ sensor.^49,50^

Moreover, our WGCNA indicated that the improvement of LVEF in rAAV5-*hS100A1* treated pigs may be due to a hitherto unknown anti-inflammatory effect of S100A1 in diseased myocardium. In support of this notion, we found an inverse correlation between LVEF improvement in our pig model and gene networks representing innate and adaptive immunity. By now, the negative impact of perpetuated inflammation in post-ischemic heart failure has been documented by a realm of studies^78,79^ and become the cornerstone of new strategies to combat cardiac dysfunction, e.g., by anti-inflammatory drugs and biologics with interleukin-1 and other inflammatory molecules as targets.^80–82^ Our systems medicine analysis corroborates the view that rAAV5-*hS100A1* therapy mitigated the activity of inflammatory gene networks in the post-MI porcine heart, which account for adverse cardiac remodeling and subsequent functional deterioration.^78,82,83^ This notion is in line with previous data showing attenuated serum levels of cytokines and chemokines by rAAV-based *S100A1* gene therapy^19^ in post-MI rodent models exhibiting, e.g., enhanced neutrophil generation and migration, such as by colony-stimulating factor 2 and 3,^84^ or convey pro-inflammatory T-cell actions, such as interferon-gamma or interleukin 17.^83,85,86^ In line with this, our WGCNA infers diminished activity of neutrophil and T-cell related gene networks in rAAV5-*hS100A1* treated post-MI pig hearts, e.g., with downregulation of cathepsin endoproteases or the CD3 T-cell receptor subunits E and D that are required for T-cell activation. In addition, as previous studies indicate a likewise detrimental contribution of B-cell mediated inflammatory processes to heart failure pathogenesis,^87^ our data further unveil diminished activity B-cell dependent gene ensembles due to rAAV5-*hS100A1* based gene therapy.

Use of complementary post-MI model in mice enabled us to further validate the novel anti-inflammatory potential of S100A1 by demonstrating mitigated expression of selected marker genes underlying neutrophil and T-cell activation in remodeled myocardium due to rAAV5-based S100A1 treatment. The administration mode of direct intramyocardial injections is likely not translatable to humans but ensured controlled delivery of the therapeutic vector system. Interestingly, recent studies showed direct binding and antagonistic effects of S100 proteins, such as S100A1, S100A4 or S100A6, on interferons and interleukins,^52^ ^53^ whose downstream signaling has been ascribed to adverse post-MI cardiac remodeling, contractile performance deterioration and mortality.^78^ Since we have previously shown that decreased S100A6 expression in endothelial cells triggered type I interferon signaling,^88^ it is tempting to speculate that rAAV5-enforced S100A1 expression may attenuate myocardial interferon signaling in our disease models. This could contribute to the overall anti-inflammatory action of S100A1 in post-MI dysfunctional myocardium, as seen in our study, and the previously reported mitigation of circulating inflammatory mediators after post-MI rAAV-based S100A1 treatment.^19^ Although the precise nature of the underlying molecular and cellular mechanisms warrants clarification in future studies, this novel observation may bear clinical relevance considering the ongoing search for new therapies to combat exaggerated inflammation that aggravates heart failure development. S100A1’s anti-inflammatory effect in the heart may also synergize with its otherwise favorable therapeutic spectrum, such as restored calcium handling and energy homeostasis, mitochondrial protection and myofilament function, as demonstrated by numerous studies by others and our group.^19–22,71^

## Conclusions and clinical perspective

Overall, our study met the aim to characterize rAAV5 as a suitable vector for cardiac-targeted gene delivery to a human-sized hearts as well as for cardioprotective gene therapy in a clinically relevant disease model by using the validated target S100A1. The introduction of various standardized quality measures for a catheter-based clinically applicable ROA and a meaningful clinical endpoint design by our CMR study enabled a first proof-of-concept study already demonstrating therapeutic efficacy of the vector-target system with a dosage ranging at the lower end of approved dose margins in current early rAAV-based CHF gene therapy trials. This corroborates the therapeutic efficacy of our GTMP candidate rAAV5*-hS100A1* and provides the rationale for further pre-clinical dose-escalating studies leveraging the full dose range of up to 3×10^14^ vgcs per patient being approved for AAV-based CHF GTMPs in ongoing FiH studies (NCT04179643). The favorable clinical safety profile of rAAV5, its high production yield and experience of regulatory bodies with this serotype, render the vector a promising candidate for expedited cardiac GTMP developmental programs using S100A1 as a therapeutic target.

## Study limitations

We used the fewest animals possible to test our hypothesis aligning with our responsibility and recommendations to limit the use of animals in biomedical research. The results and effect sizes of this study will inform subsequent IND-enabling pre-clinical studies with predefined endpoints. Our study is further limited by the sole use of male animals. Informed by our data, further IND-enabling work probing the dose-dependent safety and efficacy of rAAV5-*hS100A1* as well as the prevention of germ line transmission will require two sexes. In addition, only animals free of neutralizing serum factors for the AAVs used were enrolled to exclude confounding factors. As the amount of anti-rAAV5 seronegative animals in our local farm pig populations ranged from 60-70% (data not shown), similar to human data, subsequent pre-clinical studies may be able to exclude only a small fraction of animals from enrollment.

## Supporting information

Supplement AAV5-S100A1

## Abbreviations

AAV: Adeno-associated virus
CMR: Cardiac magnetic resonance
CRID: Catheter-based retrograde cardiac-targeted intravenous delivery
CUPID: Calcium upregulation by percutaneous administration of gene therapy in patients with cardiac diseases
CHF: Congestive heart failure
FAEs: Fatal adverse events
GTMP: Gene therapy medicinal product
GFP: Green fluorescent protein
IND: Investigational new drug
LGE: Late gadolinium enhancement
LAD: Left anterior descending artery
LCX: Left circumflex artery
LVEF: Left ventricular ejection fraction
luc: Luciferase
ME: Module eigengene
MI: Myocardial infarction
MLCv2: Myosin light chain ventricle-2
ROA: Route of administration
SAEs: Serious adverse events
vgcs: Vector genome copies
WGCNA: Weighted gene co-expression network analysis

## Acknowledgement

This study was supported by grants from the Ministry of Science, Research and the Arts of Baden-Württemberg (*forum gesundheitsstandort BW*, Translation program, subproject 4, 32-5400/58/3 to PM), German Center for Cardiovascular Research (DZHK; project 81Z0500101 to PM), National Institute of Heart, Lung and Blood Research (NHLBI; RO1 HL92139 to PM, RO1 HL0616190 and P01 HL147841 to WJK).

## Author Contributions

DK, BK, KV, JB, PS performed large animal experiments. JS, JR and FA contributed to MRI experiments. DK, EG and KP performed molecular analysis and performed small animal experiments. AJ was responsible for vector production. MB performed bioinformatic analysis. WJK, PM, DK, BK designed experiments and WJK, PM and HP supervised all studies. HK and NF provided institutional support and feedback on the manuscript. PM, DR, JR wrote and edited the manuscript. All authors approved the final version of this paper.

## References

1. Pleger ST, Brinks H, Ritterhoff J et al. Heart failure gene therapy: the path to clinical practice. Circ Res. 2013;113:792–809.

2. Hajjar RJ, Ishikawa K. Introducing Genes to the Heart: All About Delivery. Circ Res. 2017;120:33–35.

3. Kieserman JM, Myers VD, Dubey P, Cheung JY, Feldman AM. Current Landscape of Heart Failure Gene Therapy. J Am Heart Assoc. 2019;8:e012239.

4. Greenberg B, Butler J, Felker GM et al. Calcium upregulation by percutaneous administration of gene therapy in patients with cardiac disease (CUPID 2): a randomised, multinational, double-blind, placebo-controlled, phase 2b trial. Lancet. 2016;387:1178–86.

5. Hulot JS, Salem JE, Redheuil A et al. Effect of intracoronary administration of AAV1/SERCA2a on ventricular remodelling in patients with advanced systolic heart failure: results from the AGENT-HF randomized phase 2 trial. Eur J Heart Fail. 2017;19:1534–1541.

6. Pleger ST, Raake P, Katus HA, Most P. Cardiac calcium handling on trial: targeting the failing cardiomyocyte signalosome. Circ Res. 2014;114:12–4.

7. Yla-Herttuala S. Gene Therapy for Heart Failure: Back to the Bench. Mol Ther. 2015;23:1551–2.

8. Russell S, Bennett J, Wellman JA et al. Efficacy and safety of voretigene neparvovec (AAV2-hRPE65v2) in patients with RPE65-mediated inherited retinal dystrophy: a randomised, controlled, open-label, phase 3 trial. Lancet. 2017;390:849–860.

9. Schwartz M, Likhite S, Meyer K. Onasemnogene abeparvovec-xioi: a gene replacement strategy for the treatment of infants diagnosed with spinal muscular atrophy. Drugs Today (Barc). 2021;57:387–399.

10. Keam SJ. Eladocagene Exuparvovec: First Approval. Drugs. 2022;82:1427–1432.

11. Blair HA. Valoctocogene Roxaparvovec: First Approval. Drugs. 2022;82:1505–1510.

12. Marks P. Enhancing gene therapy regulatory interactions. Expert Opin Biol Ther. 2022;22:1073–1074.

13. Chhina M, Drago D, Ndu A. Planning for progress: A US regulatory approach to advancing the clinical development of gene therapies. Mol Ther. 2022;30:2397–2400.

14. Yang TY, Braun M, Lembke W et al. Immunogenicity assessment of AAV-based gene therapies: An IQ consortium industry white paper. Mol Ther Methods Clin Dev. 2022;26:471–494.

15. Philippidis A. Food and Drug Administration Lifts Clinical Hold on Pfizer Duchenne Muscular Dystrophy Gene Therapy Linked to Patient Death. Hum Gene Ther. 2022;33:573–576.

16. Philippidis A. Novartis Confirms Deaths of Two Patients Treated with Gene Therapy Zolgensma. Hum Gene Ther. 2022;33:842–844.

17. Weber C, Neacsu I, Krautz B et al. Therapeutic safety of high myocardial expression levels of the molecular inotrope S100A1 in a preclinical heart failure model. Gene Ther. 2014;21:131–8.

18. Pleger ST, Most P, Boucher M et al. Stable myocardial-specific AAV6-S100A1 gene therapy results in chronic functional heart failure rescue. Circulation. 2007;115:2506–15.

19. Katz MG, Gubara SM, Hadas Y et al. Effects of genetic transfection on calcium cycling pathways mediated by double-stranded adeno-associated virus in postinfarction remodeling. J Thorac Cardiovasc Surg. 2020;159:1809–1819 e3.

20. Pleger ST, Shan C, Ksienzyk J et al. Cardiac AAV9-S100A1 gene therapy rescues post-ischemic heart failure in a preclinical large animal model. Sci Transl Med. 2011;3:92ra64.

21. Brinks H, Rohde D, Voelkers M et al. S100A1 genetically targeted therapy reverses dysfunction of human failing cardiomyocytes. J Am Coll Cardiol. 2011;58:966–73.

22. Jungi S, Fu X, Segiser A et al. Enhanced Cardiac S100A1 Expression Improves Recovery from Global Ischemia-Reperfusion Injury. J Cardiovasc Transl Res. 2018;11:236–245.

23. Ritterhoff J, Volkers M, Seitz A et al. S100A1 DNA-based Inotropic Therapy Protects Against Proarrhythmogenic Ryanodine Receptor 2 Dysfunction. Mol Ther. 2015;23:1320–1330.

24. Fukushima H, Chung CS, Granzier H. Titin-isoform dependence of titin-actin interaction and its regulation by S100A1/Ca2+ in skinned myocardium. J Biomed Biotechnol. 2010;2010:727239.

25. Boerries M, Most P, Gledhill JR et al. Ca2+-dependent interaction of S100A1 with F1-ATPase leads to an increased ATP content in cardiomyocytes. Mol Cell Biol. 2007;27:4365–73.

26. Kettlewell S, Most P, Currie S, Koch WJ, Smith GL. S100A1 increases the gain of excitation-contraction coupling in isolated rabbit ventricular cardiomyocytes. J Mol Cell Cardiol. 2005;39:900–10.

27. Kiewitz R, Acklin C, Schafer BW et al. Ca2+-dependent interaction of S100A1 with the sarcoplasmic reticulum Ca2+-ATPase2a and phospholamban in the human heart. Biochem Biophys Res Commun. 2003;306:550–7.

28. Most P, Remppis A, Pleger ST et al. Transgenic overexpression of the Ca2+-binding protein S100A1 in the heart leads to increased in vivo myocardial contractile performance. J Biol Chem. 2003;278:33809–17.

29. Du XJ, Cole TJ, Tenis N et al. Impaired cardiac contractility response to hemodynamic stress in S100A1-deficient mice. Mol Cell Biol. 2002;22:2821–9.

30. Maco B, Mandinova A, Durrenberger MB, Schafer BW, Uhrik B, Heizmann CW. Ultrastructural distribution of the S100A1 Ca2+-binding protein in the human heart. Physiol Res. 2001;50:567–74.

31. Most P, Bernotat J, Ehlermann P et al. S100A1: a regulator of myocardial contractility. Proc Natl Acad Sci U S A. 2001;98:13889–94.

32. Yamasaki R, Berri M, Wu Y et al. Titin-actin interaction in mouse myocardium: passive tension modulation and its regulation by calcium/S100A1. Biophys J. 2001;81:2297–313.

33. Remppis A, Most P, Loffler E et al. The small EF-hand Ca2+ binding protein S100A1 increases contractility and Ca2+ cycling in rat cardiac myocytes. Basic Res Cardiol. 2002;97 Suppl 1:I56–62.

34. Most P, Seifert H, Gao E et al. Cardiac S100A1 protein levels determine contractile performance and propensity toward heart failure after myocardial infarction. Circulation. 2006;114:1258–68.

35. Raake PW, Schlegel P, Ksienzyk J et al. AAV6.betaARKct cardiac gene therapy ameliorates cardiac function and normalizes the catecholaminergic axis in a clinically relevant large animal heart failure model. Eur Heart J. 2013;34:1437–47.

36. Raake PWJ, Barthelmes J, Krautz B et al. Comprehensive cardiac phenotyping in large animals: comparison of pressure-volume analysis and cardiac magnetic resonance imaging in pig post-myocardial infarction systolic heart failure. Int J Cardiovasc Imaging. 2019;35:1691–1699.

37. aus dem Siepen F, Buss SJ, Messroghli D, et al. T1 mapping in dilated cardiomyopathy with cardiac magnetic resonance: quantification of diffuse myocardial fibrosis and comparison with endomyocardial biopsy. Eur Heart J Cardiovasc Imaging. 2015;16:210–6.

38. Riffel JH, Schmucker K, Andre F et al. Cardiovascular magnetic resonance of cardiac morphology and function: impact of different strategies of contour drawing and indexing. Clin Res Cardiol. 2019;108:411–429.

39. Muller T, Boileau E, Talyan S et al. Updated and enhanced pig cardiac transcriptome based on long-read RNA sequencing and proteomics. J Mol Cell Cardiol. 2021;150:23–31.

40. Jakobi T, Siede D, Eschenbach J et al. Deep Characterization of Circular RNAs from Human Cardiovascular Cell Models and Cardiac Tissue. Cells. 2020;9.

41. Langfelder P, Horvath S. WGCNA: an R package for weighted correlation network analysis. BMC Bioinformatics. 2008;9:559.

42. Gao E, Lei YH, Shang X et al. A novel and efficient model of coronary artery ligation and myocardial infarction in the mouse. Circ Res. 2010;107:1445–53.

43. Kupatt C, Dessy C, Hinkel R et al. Heat shock protein 90 transfection reduces ischemia-reperfusion-induced myocardial dysfunction via reciprocal endothelial NO synthase serine 1177 phosphorylation and threonine 495 dephosphorylation. Arterioscler Thromb Vasc Biol. 2004;24:1435–41.

44. Ziegler T, Bahr A, Howe A et al. Tbeta4 Increases Neovascularization and Cardiac Function in Chronic Myocardial Ischemia of Normo- and Hypercholesterolemic Pigs. Mol Ther. 2018;26:1706–1714.

45. Myers VD, Landesberg GP, Bologna ML, Semigran MJ, Feldman AM. Cardiac Transduction in Mini-Pigs After Low-Dose Retrograde Coronary Sinus Infusion of AAV9-BAG3: A Pilot Study. JACC: Basic to Translational Science. 2022;7:951–953.

46. Ibanez B, Aletras AH, Arai AE et al. Cardiac MRI Endpoints in Myocardial Infarction Experimental and Clinical Trials: JACC Scientific Expert Panel. J Am Coll Cardiol. 2019;74:238–256.

47. Puntmann VO, Carr-White G, Jabbour A et al. T1-Mapping and Outcome in Nonischemic Cardiomyopathy: All-Cause Mortality and Heart Failure. JACC Cardiovasc Imaging. 2016;9:40–50.

48. Klem I, Shah DJ, White RD et al. Prognostic value of routine cardiac magnetic resonance assessment of left ventricular ejection fraction and myocardial damage: an international, multicenter study. Circ Cardiovasc Imaging. 2011;4:610–9.

49. Duarte-Costa S, Castro-Ferreira R, Neves JS, Leite-Moreira AF. S100A1: a major player in cardiovascular performance. Physiol Res. 2014;63:669–81.

50. Ritterhoff J, Most P. Targeting S100A1 in heart failure. Gene Ther. 2012;19:613–21.

51. Takagawa J, Zhang Y, Wong ML et al. Myocardial infarct size measurement in the mouse chronic infarction model: comparison of area- and length-based approaches. J Appl Physiol (1985). 2007;102:2104–11.

52. Kazakov AS, Sokolov AS, Permyakova ME et al. Specific cytokines of interleukin-6 family interact with S100 proteins. Cell Calcium. 2022;101:102520.

53. Kazakov AS, Sofin AD, Avkhacheva NV et al. Interferon Beta Activity Is Modulated via Binding of Specific S100 Proteins. Int J Mol Sci. 2020;21.

54. Von Drygalski A, Giermasz A, Castaman G, et al. Etranacogene dezaparvovec (AMT-061 phase 2b): normal/near normal FIX activity and bleed cessation in hemophilia B. Blood Adv. 2019;3(21):3241–3247. Blood Adv. 2020;4:3668.

55. Joshi PRH, Cervera L, Ahmed I et al. Achieving High-Yield Production of Functional AAV5 Gene Delivery Vectors via Fedbatch in an Insect Cell-One Baculovirus System. Mol Ther Methods Clin Dev. 2019;13:279–289.

56. Xanthopoulos A, Starling RC, Kitai T, Triposkiadis F. Heart Failure and Liver Disease: Cardiohepatic Interactions. JACC Heart Fail. 2019;7:87–97.

57. Laribi S, Mebazaa A. Cardiohepatic syndrome: liver injury in decompensated heart failure. Curr Heart Fail Rep. 2014;11:236–40.

58. Raake PW, Hinkel R, Muller S et al. Cardio-specific long-term gene expression in a porcine model after selective pressure-regulated retroinfusion of adeno-associated viral (AAV) vectors. Gene Ther. 2008;15:12–7.

59. Gray SJ, Matagne V, Bachaboina L, Yadav S, Ojeda SR, Samulski RJ. Preclinical differences of intravascular AAV9 delivery to neurons and glia: a comparative study of adult mice and nonhuman primates. Mol Ther. 2011;19:1058–69.

60. Zincarelli C, Soltys S, Rengo G, Rabinowitz JE. Analysis of AAV serotypes 1-9 mediated gene expression and tropism in mice after systemic injection. Mol Ther. 2008;16:1073–80.

61. Pillay S, Meyer NL, Puschnik AS et al. An essential receptor for adeno-associated virus infection. Nature. 2016;530:108–12.

62. Large EE, Silveria MA, Zane GM, Weerakoon O, Chapman MS. Adeno-Associated Virus (AAV) Gene Delivery: Dissecting Molecular Interactions upon Cell Entry. Viruses. 2021;13.

63. Meyer NL, Chapman MS. Adeno-associated virus (AAV) cell entry: structural insights. Trends Microbiol. 2022;30:432–451.

64. Mendell JR, Connolly AM, Lehman KJ et al. Testing preexisting antibodies prior to AAV gene transfer therapy: rationale, lessons and future considerations. Mol Ther Methods Clin Dev. 2022;25:74–83.

65. Weber T. Anti-AAV Antibodies in AAV Gene Therapy: Current Challenges and Possible Solutions. Front Immunol. 2021;12:658399.

66. Fargnoli AS, Katz MG, Williams RD, Kendle AP, Steuerwald N, Bridges CR. Liquid jet delivery method featuring S100A1 gene therapy in the rodent model following acute myocardial infarction. Gene Ther. 2016;23:151–7.

67. Jackson BM, Gorman JH, Moainie SL et al. Extension of borderzone myocardium in postinfarction dilated cardiomyopathy. J Am Coll Cardiol. 2002;40:1160–7; discussion 1168-71.

68. Leong CN, Dokos S, Andriyana A et al. The role of end-diastolic myocardial fibre stretch on infarct extension. Int J Numer Method Biomed Eng. 2020;36:e3291.

69. Zhang Z, Sun K, Saloner DA et al. The benefit of enhanced contractility in the infarct borderzone: a virtual experiment. Front Physiol. 2012;3:86.

70. Wenk JF, Klepach D, Lee LC et al. First evidence of depressed contractility in the border zone of a human myocardial infarction. Ann Thorac Surg. 2012;93:1188–93.

71. Most P, Pleger ST, Volkers M et al. Cardiac adenoviral S100A1 gene delivery rescues failing myocardium. J Clin Invest. 2004;114:1550–63.

72. Muller JE, Rude RE, Braunwald E et al. Myocardial infarct extension: occurrence, outcome, and risk factors in the Multicenter Investigation of Limitation of Infarct Size. Ann Intern Med. 1988;108:1–6.

73. Chopra N, Yang T, Asghari P et al. Ablation of triadin causes loss of cardiac Ca2+ release units, impaired excitation-contraction coupling, and cardiac arrhythmias. Proc Natl Acad Sci U S A. 2009;106:7636–41.

74. Quiros PM, Prado MA, Zamboni N et al. Multi-omics analysis identifies ATF4 as a key regulator of the mitochondrial stress response in mammals. J Cell Biol. 2017;216:2027–2045.

75. House SL, Bolte C, Zhou M et al. Cardiac-specific overexpression of fibroblast growth factor-2 protects against myocardial dysfunction and infarction in a murine model of low-flow ischemia. Circulation. 2003;108:3140–8.

76. Seo HR, Jeong HE, Joo HJ et al. Intrinsic FGF2 and FGF5 promotes angiogenesis of human aortic endothelial cells in 3D microfluidic angiogenesis system. Sci Rep. 2016;6:28832.

77. Yang N, Parker LE, Yu J et al. Cardiac retinoic acid levels decline in heart failure. JCI Insight. 2021;6.

78. Westman PC, Lipinski MJ, Luger D et al. Inflammation as a Driver of Adverse Left Ventricular Remodeling After Acute Myocardial Infarction. J Am Coll Cardiol. 2016;67:2050–60.

79. Blanton RM, Carrillo-Salinas FJ, Alcaide P. T-cell recruitment to the heart: friendly guests or unwelcome visitors? Am J Physiol Heart Circ Physiol. 2019;317:H124–H140.

80. Hanna A, Frangogiannis NG. Inflammatory Cytokines and Chemokines as Therapeutic Targets in Heart Failure. Cardiovasc Drugs Ther. 2020;34:849–863.

81. Abbate A, Toldo S, Marchetti C, Kron J, Van Tassell BW, Dinarello CA. Interleukin-1 and the Inflammasome as Therapeutic Targets in Cardiovascular Disease. Circ Res. 2020;126:1260–1280.

82. Libby P. Targeting Inflammatory Pathways in Cardiovascular Disease: The Inflammasome, Interleukin-1, Interleukin-6 and Beyond. Cells. 2021;10.

83. Bansal SS, Ismahil MA, Goel M et al. Activated T Lymphocytes are Essential Drivers of Pathological Remodeling in Ischemic Heart Failure. Circ Heart Fail. 2017;10:e003688.

84. Tecchio C, Micheletti A, Cassatella MA. Neutrophil-derived cytokines: facts beyond expression. Front Immunol. 2014;5:508.

85. Mora-Ruiz MD, Blanco-Favela F, Chavez Rueda AK, Legorreta-Haquet MV, Chavez-Sanchez L. Role of interleukin-17 in acute myocardial infarction. Mol Immunol. 2019;107:71–78.

86. Levick SP, Goldspink PH. Could interferon-gamma be a therapeutic target for treating heart failure? Heart Fail Rev. 2014;19:227–36.

87. Garcia-Rivas G, Castillo EC, Gonzalez-Gil AM, et al. The role of B cells in heart failure and implications for future immunomodulatory treatment strategies. ESC Heart Fail. 2020;7:1387–1399.

88. Lerchenmuller C, Heissenberg J, Damilano F et al. S100A6 Regulates Endothelial Cell Cycle Progression by Attenuating Antiproliferative Signal Transducers and Activators of Transcription 1 Signaling. Arterioscler Thromb Vasc Biol. 2016;36:1854–67.

