## Supplement AAV5-S100A1 for "Cardiac-targeted rAAV5-S100A1 gene therapy protects against adverse remodeling and contractile dysfunction in post-ischemic hearts"

- **Appendix: Extended Methods**
- **Appendix: Extended Figures**

### Appendix: Extended methods:

All animal procedures and experiments were carried out according to the 'Guide for the Care and Use of Laboratory Animals' (National Institutes of Health) and were approved by the local Institutional Animal Care and Use Committee of Baden-Württemberg, Germany and Jefferson University, PH, USA.

All mice were housed at 22°C with a 12-hour light, 12-hour dark cycle with free access to water and standard chow. Bay cage types are used for housing of pigs. Per pig, a body weight prescribed floor space of at least 0.5 and 0.7 $m^2$ were provided according to directive 2010/63/EU with straw bedding. The environment was enriched by pellet balls, chains and gnawing rods. Humidity and temperature were kept at 50-60 % and 20-24 °C, respectively. Access to water was unlimited and restricted food provided twice/d (SAF130M).

#### rAAV manufacturing

rAAV vectors were produced, purified, titrated, and stored as described previously.[1-3] In more detail, high titre vectors were produced with a double transfection approach of 293T cells in 10-layer cell stacks. For production of rAAV5-*human S100A1 (hS100A1)*, rAAV5-*luc*, and rAAV5-*green fluorescent protein (gfp) and r*AAV5-*hS100A1-IRES-gfp* vectors, pDP5rs providing the rAAV-5 cap sequence as well as the adenoviral helper sequences were co-transfected with pdsCMV-MLC0.26-*hS100A1*, pdsCMV-MLC0.26-*luc* and pdsCMV-

MLC0.26-*gfp* and pdsCMV-MLC0.26-*hS100A1-IRES-gfp* respectively, while CMV refers to a short sequence of the full length CMV promoter to boost expression of the cardiac-biased MLC0.26 expression fragment. The sequence of the utilized *hs100A1* cDNA and production of rAAV6-*luc* and rAAV9-*luc* with the CMV-MLC0.26 expression cassette has been published elsewhere.[1,2] Viral vectors were harvested from the cells as well as from the supernatant after 48h-72h. Freeze and thaw treated cell lysates and ammonium-sulfate precipitated supernatants were purified by Iodixanol step gradients. 20ml of cell lysate and supernatant were layered onto a 4-phase iodixanol gradient (15%, 25%, 40% and 60%), respectively. Viral genome copy (vgc) numbers were quantified using qPCR. A plasmid standard curve was built from 1:10 serial dilutions. Before setting up the qPCR mix the viral genome was released from the viral capsids via an alkaline lysis by incubating 10µl of the viral vector with 10µl of Tris-EDTA and 20µl of 2M NaOH by 56°C. This reaction was neutralized after 30min by 960µl of 40mM HCl. The qPCR mix was prepared using the QuantiTect SYBR Green PCR + UNG Kit (Qiagen, Hilden, Germany) and pipetted in triplicates. A CFX96 Touch Real Time PCR detection system device (Bio-Rad, München, Germany) was used with a thermal protocol of 2 min / 50 °C, 15 min / 94 °C, followed by 40 cycles of 15 s / 94 °C, 30 s / 60 °C and 30 s / 72 °C, ended with the generation of a melting curve (increase of temperature from 50 to 95 °C with 0.5 °C per 5 s). Samples that were either reported undetectable or detected ≥ Cq 35 were not counted. Vgc were calculated by means of a non-linear regression analysis using Excel and GraphPad

**Cardiac-targeted gene delivery to porcine hearts**

Cardiac-targeted catheter-based retrograde intravenous delivery (CRID) of AAV vectors both to normal and infarcted pig hearts was carried out as described elsewhere.[1-3] To this end, male neutered German farm pigs (mean body weight: 31.5 ± 1.85 kg) were anesthetized with an intramuscular injection of ketamine 15 mg/kg body weight (Ketamin 10 %, Betapharm, Augsburg, Germany) and midazolam 1 mg/kg body weight (Hoffmann-La Roche, Grenzach-Wyhlen, Germany). Periprocedural analgesia 0.3 mg buprenorphine (Buprenovet multidose

0.3 mg/ml, Bad Homburg vor der Höhe, Germany) was administered through an IV catheter in the ear vein. After intubation anesthesia was maintained by inhalation of 2 % isoflurane (Isofluran CP 1 ml/ml, CP-Pharma, Burgdorf, Germany). Then two catheter introducer sheaths were placed using a 7F for the right carotid artery and an 8F for the right external jugular vein. Full anticoagulation was achieved by i.v. application of 5000 IU per hour of heparin (Heparin-Natrium LEO 25.000 I.E./5 ml, Leo Pharma, Ballerup, Denmark). A retroinfusion catheter (Attain Clarity 6225, Medtronic, Dublin, Ireland) was placed in the anterior intraventricular vein (AIV) while a 6 F guiding catheter (6 F Judkins right, Cordis, Cardinal Health, Dublin, Ireland) was placed in the left coronary artery. A PTCA balloon catheter (Emerge 3.5 x 15 mm, Boston Scientific, Natick, US) was placed via a 0.014" guiding wire (Balance Middle Weight, Abbott, Santa Clara, US) in the LAD distal to the first diagonal branch. The LAD was temporarily occluded by that balloon during retrograde delivery of the AAV vectors. For the comparative biodistribution analysis, AAV5-*luc*, AAV6-*luc* and AAV9-*luc* were each applied with a single dosage of $1x10^{13}$ vgcs to n=5 animals per group. To ensure a standardized CRID protocol, procedural checkpoints were implemented: first, appropriate positioning of the retroinfusion catheter in the cardiac anterior interventricular vein (AIV), closely to the outflow of the coronary sinus, was recorded by fluoroscopic images. Concomitantly, the adequate position of the intracoronary balloon catheter was ascertained to avert an immediate antegrade flushing of the myocardial capillary system during retrograde AAV installation (appendix figure A1). Then, contrast agent was retrogradely administered over the blocked venous catheter to rule out an immediate leakage into the systemic circulation and avoid selective transfer into smaller venous branches of the AIV during rAAV infusion (appendix figure A2). Once these checkpoints were met in each animal, vectors were applied by CRID and biodistribution was systematically assessed after 30 days by luminometric assessment of *luc* reporter gene activity in porcine heart homogenates and additional organ samples. For the therapy study, infarcted animals were randomized either to post-MI AAV5-*hS100A1* (n=5) treatment or post-MI control procedure (n=4; 3 x AAV5-hRluc and 1 x saline) with a vector dosage of $1\times10^{13}$. For the administration, AAV5 vectors were diluted in 48 ml of 0.9 % saline solution. The total amount was divided in three portions of 16 ml. During the injection the two PTCA balloons (one in the LAD, one at the tip of the retroinfusion catheter in the AIV were inflated to block venous outflow and arterial inflow. The balloon in the AIV was inflated for 3 min while the balloon in the LAD was inflated for 45 sec each time. Before each portion of vector solution 2 ml of 1:10 diluted nitroglycerine (Nitrolingual infus. 1 mg/ml, Pohl-Boskamp GmbH, Hohenlockstedt, Germany) were administered in the AIV. Between the repeated injections both balloons were deflated for 2 min to allow reflow. At the beginning and end of the procedure an angiography was performed to check the correct position of the catheters and the integrity of the LAD and AIV.

**Tissue vector abundance, gene expression and immunoblot analyses**

Isolation of tissue DNA and RNA including reverse transcription and PCR-based vector genome abundance, gene expression quantification as well as immunoblotting was conducted as described previously.[1-3] For biodistribution and expression analyses, genomic DNA and total RNA from heart as well as from liver, lung, kidney and brain samples were isolated and analyzed via quantitative real-time PCR (qPCR).

*Isolation of gDNA and total RNA:* Tissue pieces were homogenized in a two-step procedure. After adding 8 µl/mg ice-cold lysis buffer (1 x DPBS without $Ca^{2+}$/$Mg^{2+}$ (Merck, Darmstadt, Germany) and 10 mM EDTA pH 8.0 (Merck, Darmstadt, Germany)) tissue pieces were treated using TissueRuptor II system (Qiagen, Hilden, Germany) for a first round of homogenization. Aliquots were stored at -80 °C or subjected directly to gDNA and total RNA extraction using the AllPrep DNA/RNA Mini Kit (Qiagen, Hilden, Germany) with a slightly modified protocol. Briefly, 200-300 µl tissue homogenates were mixed with 600 µl RLT Plus buffer and treated with the PreCellys 24 (VWR, Darmstadt, Germany) bead mill system at 5.500 $min^{-1}$ for 30 s using 2.8 mm beads for a second round of homogenization. The RLT Plus homogenates were then centrifuged for 3 min at 18.000 x g and the supernatants were transferred to AllPrep DNA spin columns. Genomic DNA and total RNA were purified by following steps 4 to 16 from the manufacturer's handbook (version October 2019) and eluted in RNase-free water. To avoid genomic DNA contamination of the total RNA preparations, an additional on-column DNAse digestion was carried out by replacing step 8 with steps E1-E4 from the manufacturer's handbook (RNase-free DNase Set, Qiagen, Hilden, Germany). Nucleic acid concentrations were determined photometrically using a spectrophotometer (NanoDrop 2000, Thermo Fisher Scientific, Waltham, Germany). Samples with a total RNA concentration lower than 65 ng/µl were concentrated for 10 min at 45 °C in the Concentrator plus (Eppendorf, Hamburg, Germany) up to two times. In case of poor gDNA results, extraction was repeated with the DNeasy Blood and Tissue Kit (Qiagen, Hilden, Germany). For this purpose, 200 µl tissue homogenate were incubated with 20 µl proteinase K over night at room temperature. After having followed steps 2 to 7 from the manufacturer's handbook (version July 2020) gDNA was eluted in RNase-free water. gDNA was stored at +4 °C for short term, resp. -20 °C for long term, total RNA was stored at -80 °C if it was not used immediately for cDNA synthesis. *cDNA-synthesis:* Total RNA was reverse transcribed to cDNA using the QuantiTect Reverse Transcription Kit (Qiagen, Hilden, Germany) according to the manufacturer's handbook (version March 2009).

*Detection of viral genome copies (vgc) of S100A1 and hRluc in isolated porcine gDNA and synthetized cDNA:* Vgc were quantified both in isolated gDNA and synthetized cDNA samples with a standard curve built from 1:10 serial dilutions (steps 2 to 8) of a standardized plasmid DNA. The qPCR mix was prepared using QuantiTect SYBR Green PCR + UNG Kit (Qiagen, Hilden, Germany) and pipetted at least in duplicates. The plates were analyzed by a CFX96 Touch Real Time PCR detection system (Bio-Rad, München, Germany) with a thermal protocol of 2 min / 50 °C, 15 min / 94 °C, followed by 40 cycles of 15 s / 94 °C, 30 s / 60 °C and

30 s / 72 °C, ended with the generation of a melting curve (increase of temperature from 50 to 95 °C with 0.5 °C per 5 s). Samples that were either reported undetectable or detected ≥ Cq 35 were not counted. Vgc per µg isolated gDNA/RNA were calculated by means of a non-linear regression analysis using Excel and GraphPad. Primers (see **Fehler! Verweisquelle konnte nicht gefunden werden.**) used for detection of vgc of human S100A1 and renilla luciferase were tested for specificity beforehand using multiplex qPCR.

Table 1: Primers/Probes that were used for specificity testing in a multiplex approach, primers in bold were also used for quantification via SYBR-qPCR

| Target | Primer/probe | Sequence |
|---|---|---|
| **human s100a1** | **Forward** | **5'-TGTCCACTCCCAGTTCAATTAC-3'** |
| | **Reverse** | **5'-AACACGTTGATGAGGGTCTC-3'** |
| | Probe | 5'-*Rox*-ATCCGCGATGGGCTCTGAGCT-*TQ3*-3' |
| **hRluc** | **Forward** | **5'- TGTCCACTCCCAGTTCAATTAC -3'** |
| | **Reverse** | **5'- ATCATGCGTTTGCGTTGC -3'** |
| | Probe | 5'-HEX- ACCATGGCTTCCAAGGTGTACGAC –TQ2-3' |
| porcine casq2 | Forward | 5'-TGTGGCCTTTGCAGAGAGGA-3' |
| | Reverse | 5'-GGGTCGATCCACACGATGCT-3' |
| | Probe | 5'-*FAM*-CGCCAGGGACAACACAGACAACCCGGA-*TQ-2*-3' |

*Assessment of murine myocardial gene expression:* Total RNA isolation from LV tissue samples was performed applying the TRIZOL method, according to the manufacturer's protocol (Invitrogen) as previously described.[4] Quality of RNA was assessed by running an aliquot on a denaturating agarose (1%) gel. First strand cDNA synthesis from 1µg of total RNA was carried out by the use of the iScript cDNA Synthesis Kit (BioRad). For quantitative PCR, 5 µl of diluted cDNA (1/100) was added to a 25 µl mixture that contained a 1× concentration of iQ SYBR Green Supermix (BioRad) and 100 nM of gene-specific oligonucleotides. Subsequently, quantitative PCR was carried out on a MyiQ Single-Color Real-Time PCR detection system (BioRad) with purchased primer (Origene) for murine beta-actin (NM_007393) CAT#: MP200232, cxcr2 (NM_009909) CAT#: MP206804, Cd86 (NM_019388) CAT#: MP201956**,** Cd4 (NM_013488) CAT#: MP201927, Il1b

(NM_008361)CAT#: MP206724, TNF (NM_013693) CAT#: MP217748 and INFg (NM_008337) CAT#: MP206683. Beta-actin mRNA levels were not different between groups and used for normalization. After each run, saturation of each amplification cycle was controlled by the use of MyiQ software (version 1.0) and, subsequently a melting curve acquired by heating the product to 95°C, cooling to and maintaining at 55°C for 20 seconds, then slowly (0.5°C/s) heating to 95°C was used to determine the specificity of the PCR products, which were then confirmed by gel electrophoresis.

*Immunoblotting:* LV protein expression analysis from porcine and murine tissue was carried out as described in detail elsewhere.[1,2,4] Briefly, LV tissue samples were homogenized with a tissue raptor in lysis buffer (PBS, pH 7.4, NP40 1%, 1 mM EGTA/EDTA, protease inhibitor (1 tablet/10 ml) (Roche Applied Science; Mini Complete EDTA free protease inhibitor). Protein content was assessed by the Bio-Rad DC protein assay. Protein lysates were subjected to electrophoresis (4-20% tris-glycine gradient gels, ANAMED), transferred to a PVDF membrane (Immobilon FL, Millipore) and probed with appropriate sets of primary antibodies to assess protein levels of S100A1 (Acris, 1:1000), CSQ (Calbiochem, 1:1000), GAPDH (Millipore, 1:30000) and GFP (Clonetech, 1:30000) and fluorescent-labeled secondary antibodies (LICOR Odyssey, 1:10000). Proteins were visualized with a LI-COR infrared imager (Odyssey), and quantitative densitometric analysis was performed by applying Odyssey version 1.2 infrared imaging software.

**Pig myocardial infarction model**

Post-myocardial infarction cardiac dysfunction in male German farm pigs was induced by percutaneous transluminal temporary occlusion of the left circumflex artery (LCX) as described elsewhere.[1-3] For induction of ischemic heart dysfunction male neutered German farm pigs (mean body weight: 31.5 ± 1.85 kg) were anesthetized with an intramuscular injection of ketamine 15 mg/kg body weight (Ketamin 10 %, Betapharm, Augsburg, Germany) and midazolam 1 mg/kg body weight (Hoffmann-La Roche, Grenzach-Wyhlen, Germany). For perioperative analgesia 0.3 mg buprenorphine (Buprenovet multidose 0.3 mg/ml, Bad Homburg vor der Höhe, Germany) were administered through an IV catheter in the ear vein. After intubation anesthesia was maintained by inhalation of 2 % isoflurane (Isofluran CP 1 ml/ml, CP-Pharma, Burgdorf, Germany). Under sterile conditions a 7 F catheter introducer sheath was introduced into the right carotid artery and a 6 F diagnostic catheter (6 F Judkins right, Cordis, Cardinal Health, Dublin, Ireland) was advanced into the ostium of the left coronary artery. Full anticoagulation was achieved by i.v. application of 5000 IU per hour of heparin (Heparin-Natrium LEO 25.000 I.E./5 ml, Leo Pharma, Ballerup, Denmark). Myocardial infarction was induced by temporary (2 h) occlusion of the LCX with a percutaneous transluminal coronary angioplasty (PTCA) balloon (Emerge 3.5 x 15 mm, Boston Scientific, Natick, US) which was placed into the very proximal LCX via a 0.014" guiding wire (Balance Middle Weight, Abbott, Santa Clara, US) and inflated at 6 bar. A potential leakage was ruled out by coronary angiography. 150 mg Amiodaron (Cordarex 150 mg/3 ml, Sanofi GmbH, Frankfurt, Germany) were administered intravenously before (bolus) and during (drip infusion) LCX occlusion minimizing the occurrence of fatal arrhythmias. During the procedure animals were monitored via ECG and capnography. Two weeks (15,1 ± 0,9 d) after MI pigs were anesthetized as described above for evaluation of infarct size and cardiac function by cardiac MRI. During the examination anesthesia was maintained by continuous intravenous infusion of ketamine (3 mg/kg/h) and midazolam (0.5 mg/kg/h), boluses of propofol (Propofol 1 % MCT Fresenius Kabi AG, Bad Homburg v. d. Höhe, Germany) were given as needed. Only animals with an enhanced mass > 14 % of the left ventricle were selected for gene transfer.

**Cardiac magnetic resonance and imaging analysis**

Assessment of left ventricular (LV) ejection fraction (EF) and myocardial infarct (MI) size was performed by published standard cardiac magnetic resonance protocols adopted to porcine anatomy.[5-7] Standard CMR was performed in a right-lateral position in a 1.5 T Ingenia™ whole-body scanner (Philips Healthcare, Best, The Netherlands), with a commercial cardiac phased-array receiver coil as previously described. Following localizing scans, cine long axis 2-, 3- and 4-chamber (Ch) views as well as short axis cine (SAX) images covering the whole LV from the annulus of the atrioventricular valves to the apex (8 mm slice thickness, no gap between each slice) were obtained using a segmented-k-space balanced steady-state free precession sequence (bSSFP) employing retrospective ECG or pulse oximetric gating with 35 phases per cardiac cycles. The following imaging parameters were used: repetition time (TR) 2.8 ms; echo time (TE) 1.4 ms; flip angle (FA) 60 °. Late gadolinium enhancement (LGE) images were acquired 10 minutes after administration of 0.14 mmol/ kg body weight contrast medium (Gadobutrol (Gadovist), Schering, Berlin, Germany). LGE images were acquired employing a T1-weighted inversion recovery-prepared fast gradient echo sequence with an optimized inversion time in the same orientation as the cine images. The inversion time was adapted individually to suppress signal of normal myocardial tissue (TR 3.0 ms; TE 6 ms; FA 25 °). All images were analyzed using a commercially available semiautomatic software (CVI cmr[42], Version 5.6.6, Circle Cardiovascular Imaging Inc., Calgary, Canada). All post processing measurements were obtained as previously described. End-diastolic, end-systolic volume, ejection fraction (EF %) and LV myocardial mass (LV-M) were acquired in SAX stacks by manually tracing epi- and endocardial borders, excluding papillary muscles from the myocardium. The LV outflow tract was included as LV blood volume. Papillary muscles and trabeculae were included as myocardial tissue. LV-M = (total epicardial volume − total endocardial volume) × 1.05 g/ml. Septal wall and free wall thickness were assessed using SAX basal slice. LV end-diastolic and end-systolic diameter were obtained in 3-Ch view at the mitral chordae level basal to the tips of the papillary muscles. Regions with LGE were verified in at least one other orthogonal plane and in the same plane being obtained as a second image after changing the direction of readout. Native myocardial T1 maps were calculated from Modified Look-Locker Inversion recovery

(MOLLI, 5 s(3 s)3 s scheme) images acquired before and 15 minutes after contrast agent injection. Pre-contrast MOLLI sequence (5 s(3 s)3 s variant) used following parameters: TR 2.3 ms, TE 1.06 ms, FA 35 ° and post-contrast MOLLI sequence (4s(1 s)3 s(1s)2s) variant): TR 2.4 ms, TE 1.08 ms, FA 35 °. After visual inspection of all segments and exclusion of those segments with evidence of artefact, pre- and post-contrast T1 maps were generated using CVI cmr[42] software (Version 5.6.6, Circle Cardiovascular Imaging Inc., Calgary, Canada). Endocardial and epicardial borders were defined manually, using an offset of 10 % to avoid partial-volume effects in the subendocardial and subepicardial layers. The global T1-value was calculated as a mean of all segments with respect to the segments area. T1 values for blood were detected by manually drawing a region of interest in the LV cavity. If more than two segments in T1 showed evidence of artefacts, the images were excluded from further analysis.

**RNA-sequencing of porcine myocardium and weighted gene correlation network analysis**

Next-generation sequencing of isolated myocardial LV RNA and subsequent weighted gene correlation network analysis was carried out as described elsewhere.[8-10] Total RNA was isolated as described above. Quantity and quality were measured with Qubit (Thermo Fisher Scientific, Darmstadt, Germany) and Bioanalyzers 2100 (Agilent, Waldbronn, Germany) technology and sequencing libraries were prepared with TruSeq RiboZero Kits (Illumina, Santa Clara, CA) for sequencing with a NextSeq 550 (Illumina, Santa Clara, CA) device. Analysis of FASTQ files was performed in R (version 4.1.2).[11] FASTQ files were subjected to quality control using fastqcr (version 0.1.2)[12] and FastQC (version 0.11.8).[13] Reads were mapped against the pig genome (Sscrofa11.1_GCF_000003025.6) using Rsubread (version 2.7.1)[14] and features (mRNA, transcript IDs) were also extracted with the help of Rsubread (annotation release 106). For pathway analyses, pig annotations were mapped to corresponding human annotations using org.Ss.eg.db (version 3.13.0)[15] and org.Hs.eg.db (version 3.13.0).[16] Annotations that could not be mapped with these packages (14.9 %) were blasted with blastn from ncbi-blast+ (version 2.9.0-2)[17] with rBLAST (version 0.99.2)[18] and the best hit for mapping was identified as subject hit with sequence identity above 80 % and the lowest mean e-value below 0.01 and assigned to the queried annotation. In total, 93.15 % of all annotations could be mapped from pig to human annotations. Gene expression was then analyzed with DESeq2 (version 1.33.1)[19] prior to weighted correlation network analysis (WGCNA, version 1.70-3, [-11])[20] and additional analyses with main packages pcaExplorer (version 2.19.0)[21,22], PCAtools (version 2.5.0),[23] clusterProfiler (version 4.1.0)[24,25] and ReactomePA (version 1.37.0)[26] based on variance stabilizing transformation (VST)-normalized feature counts. For WGCNA signed network setting was used assuming scale free topology at power β of 20. Tree cutting was performed with default values of the hybrid system to combine dendrogram cutting and partitioning around medoids (PAM). Deep split was performed using the default mean value 2.[27,28]

**Mouse myocardial infarction model and intramyocardial injections**

Induction of myocardial infarction by experimental ligation of the left anterior descending (LAD) artery and intramyocardial injections of AAV5 vectors were performed as described previously.[4,29] Briefly, under general anesthesia with isoflurane (2%), the heart was exposed via a small left thoracotomy. For intramyocardial injections on both sides of the LAD, a sterile Hamilton syringe with a 30-gauge sterile beveled needle was used. In the rAAV5-*hS100A1* and rAAV5-*gfp* groups, at total dosage of $2\times10^{11}$ vgcs per animal was quickly applied by two intramyocardial injections per side with a volume of 2 µl each and a suture ligation was subsequently placed at the distal 1/2 of the LAD with 6.0 silk suture. Upon ligation, the heart was immediately placed back to the intrathoracic space followed cy closure of the skin suture and manual evacuation of pneumothoraces. Sham intramyocardial injections (saline) and operation, respectively was performed in a same manner except that the LAD was left unligated. After MI, the animals remained in a supervised setting until fully conscious. Early surgeryrelated death assigned to severe thoracic bleeding during and less than 6 hours after surgery was similar in all groups (10 %) and excluded from final analysis. MI size was quantified as previously reported.[30]

**Mouse echocardiography**

Transthoracic two-dimensional echocardiography (TTE) in lightly anesthetized mice (tribromoethanol/amylene hydrate; Avertin; 2.5% wt/vol, 8 µl/g IP) with spontaneous respiration was performed with a 12-MHz probe both in sham and infarcted mice as described in detail elsewhere.[4] TTE in M-mode was carried out in the parasternal short axis before and 28 days after the surgical procedure to assess LV diameter and subsequently fractional shortening (FS% = [(LVEDD -LVESD)/LVEDD) × 100]).

### Appendix: Supplemental figures

**Appendix figure A1**

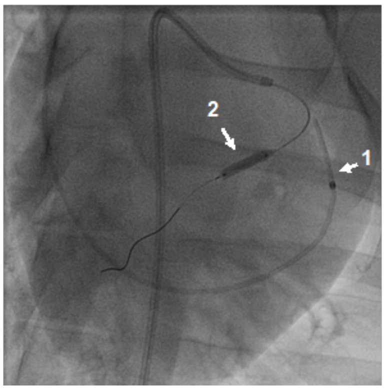

**Appendix figure A2**

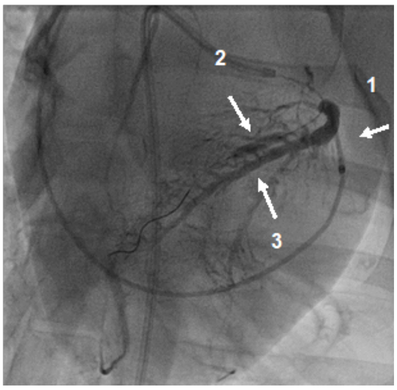

**Appendix figure A1. and A2** show representative fluoroscopic images of the standardized procedure (SOP) for cardiac-targeted catheter-based retrograde intravenous delivery (CRID) of AAVs to the porcine left ventricle as described previously.[22,25,40] Appendix 1A depicts appropriate positioning of the retroinfusion catheter in the coronary sinus with subsequent infusion into the anterior interventricular vein (AIV) (arrow 1) and a second catheter in the left anterior descending (LAD) artery (balloon inflated at the time of the image; arrow 2). Appendix 1B illustrates intravenous installation of contrast agent through the retroperfusion catheter shortly blocked distal of the coronary sinus outflow. This enables retrograde perfusion of the smaller venous branches along the entire length of the AIV and thereby later optimal access of AAVs to the left ventricle (LV). The retrograde perfusion test prior to AAV administration further excludes selective infusion of AAVs into smaller venous AIV branches that may entail poor gene transfer to the LV and, at the same time, avoids leckage through inappropiate blocking of the retroperfusion catheter that may result in systemic spilling of AAV during CRID. Brief concomittant blockade of LAD perfusion by the intracoronary catheter prevents premature wash-out of the installed virus as previously reported.

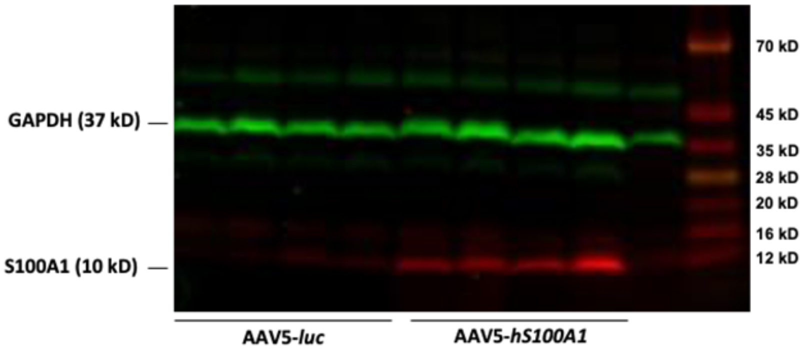

**Appendix figure A3** shows a representative immunoblot of rAAV5-*luc* and rAAV5-*hS100A1* treated post-MI LV homogenates probed with an anti-GAPDH, anti-S100A1 antibody and appropiate secondary antibodies for near-infrared fluorescent imaging. GAPDH (green signal) shows comparable loading across all samples while anti-S100A1 staining (red signal) documents robust overexpression of S100A1 protein in rAAV5-*hS100A1* over *r*AAV5-*luc* treated hearts 30 days after CRID. n=4 animals per group.

**Appendix figure B1** **Appendix figure B2** **Appendix figure B3**

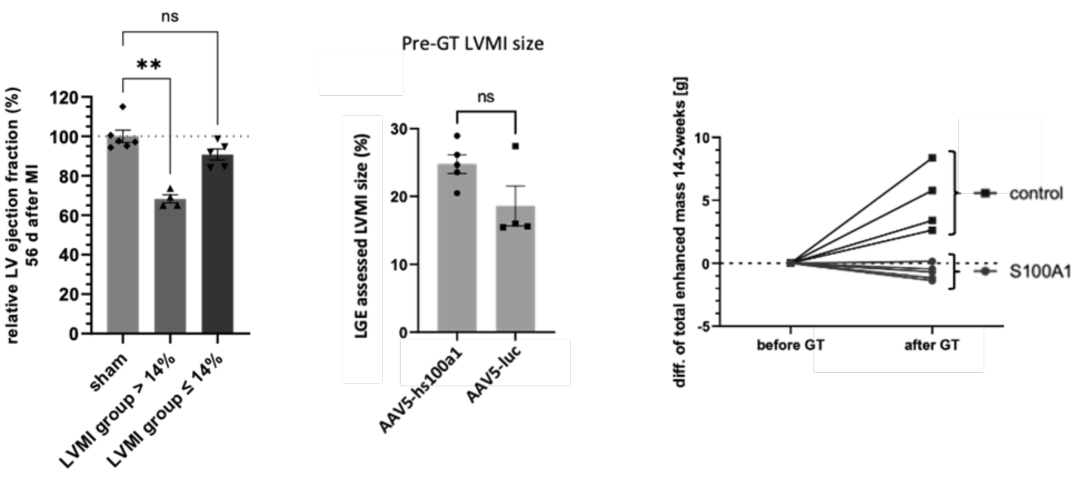

**Appendix figure B1** illustrates that a significant decline in LVEF occured in post-MI pigs with an experimental LVMI size exceeding 14% of LV mass compared to sham animals as assessed by CMR LGE. LVMI group >14; n=4 animals, LVMI ≤ 14%; n=6 animals, sham; n=6 animals. **P<0.003 sham vs. LVMI-group > 14%. Data are presented as mean+-SEM. **Appendix figure B2** depicts similar relative LVMI size prior to the rAAV5-based intervention assessed by LGE CMR in post-MI animals. Animals were assigned to the rAAV5-*hS100A1* (n=5) and rAAV5-*luc* (n=4) group which entailed a slightly greater but statistically not significant LVMI size in the rAAV5-*hS100a1* group compared to the rAAV5-*luc group* prior to treatment. **Appendix figure B3** shows LVMI extension in each control animal but in none of the rAAV5-*hS100A1* treated pigs that slightly trended eventowards regression of LVMI size. Statistical comparisons were conducted by Kruskal-Wallis test with Dunn's multiple comparisons test (B1) and Mann whitney test (B2).

**Appendix figure B4**

| | | EDV (ml) | ESV (ml) | HR (bpm) |
|---|---|---|---|---|
| control | Pig 17 | 127.01 | 58.15 | 55 |
| | Pig 33 | 162.62 | 58.03 | 49 |
| | Pig 100 | 148.33 | 61.24 | 84 |
| | Pig 101 | 90.9 | 41.29 | 80 |
| | n | 4 | 4 | 4 |
| | mean ± SEM | 132.2 ± 15.59 | 54.68 ± 4.52 | 67 ± 8.78 |
| hS100A1 | Pig 6 | 111.72 | 28.05 | 45 |
| | Pig 10 | | | 60 |
| | Pig 28 | 144.48 | 46.73 | 57 |
| | Pig 29 | 134.09 | 42.87 | 62 |
| | Pig 38 | 148.13 | 49.96 | 51 |
| | n | 4 | 4 | 5 |
| | mean ± SEM | 134.6 ± 8.19 | 41.9 ± 4.84 | 55 ± 3.11 |

**Appendix figure B4** summarizes further CMR-based data for control and rAAV5-hS100A1 treated animals. Note that animal 10 data for EDV and ESV could not be aquired due to motion artefacts.

Appendix figure C

| | | control | | | hS100A1 | | |
|---|---|---|---|---|---|---|---|
| | | mean | SEM | n | mean | SEM | n |
| leukocytes | /nl | 18,36 | ±1,25 | 4 | 17,19 | ±0,82 | 5 |
| erythrocytes | /pl | 6,05 | ±0,28 | 4 | 5,92 | ±0,17 | 5 |
| hemoglobin | g/dl | 10,20 | ±0,46 | 4 | 9,88 | ±0,29 | 5 |
| hematocrit | l/l | 0,32 | ±0,01 | 4 | 0,31 | ±0,01 | 5 |
| MCV | fl | 52,75 | ±1,89 | 4 | 53,20 | ±1,36 | 5 |
| MCH | pg/cell | 17,00 | ±0,58 | 4 | 16,80 | ±0,37 | 5 |
| MCHC | g/dl | 32,00 | ±0,41 | 4 | 31,40 | ±0,40 | 5 |
| RDW | % | 16,60 | ±0,35 | 4 | 16,76 | ±0,32 | 5 |
| Platelets | /nl | 435,50 | ±19,05 | 4 | 366,80 | ±51,77 | 5 |
| sodium | mmol/l | 134,25 | ±9,12 | 4 | 141,00 | ±0,41 | 4 |
| potassium | mmol/l | 4,23 | ±0,57 | 4 | 4,01 | ±0,11 | 5 |
| creatinine | mg/dl | 1,43 | ±0,17 | 4 | 1,33 | ±0,10 | 5 |
| urea | mg/dl | 26,00 | ±2,20 | 4 | 28,40 | ±1,94 | 5 |
| glucose | mg/dl | 91,50 | ±18,50 | 4 | 92,00 | ±2,00 | 5 |
| hs-troponin T | pg/ml | 5,00 | ±0,41 | 4 | 4,40 | ±0,60 | 5 |
| LDH | U/l | 536,50 | ±63,37 | 4 | 512,40 | ±35,12 | 5 |
| GOT/AST | U/l | 39,25 | ±3,88 | 4 | 39,00 | ±6,20 | 5 |
| GPT/ALT | U/l | 69,25 | ±9,58 | 4 | 71,20 | ±10,82 | 5 |
| GGT | U/l | 45,00 | ±11,97 | 4 | 40,60 | ±2,32 | 5 |
| lipase | U/l | 14,00 | ±1,08 | 4 | 13,00 | ±0,00 | 4 |
| albumin | g/l | 31,80 | ±2,40 | 4 | 32,64 | ±1,59 | 5 |
| Quick | % | 94,53 | ±4,79 | 3 | 91,82 | ±4,60 | 5 |
| aPTT | s | 10,03 | ±0,13 | 3 | 9,90 | ±0,15 | 4 |

**Appendix figure C** summarizes blood count and clinical chemistry parameters for control and rAAV5-*hs100a1* treated animals 70 days post-GT. This time point prior to the final functional and molecular assessments 3-month post-GT was chosen to avoid an impact of any sort of experimental manipulation on blood count or clinical chemistry. None of the tested parameters indicated abnormal alterations.

**Appendix figure D**

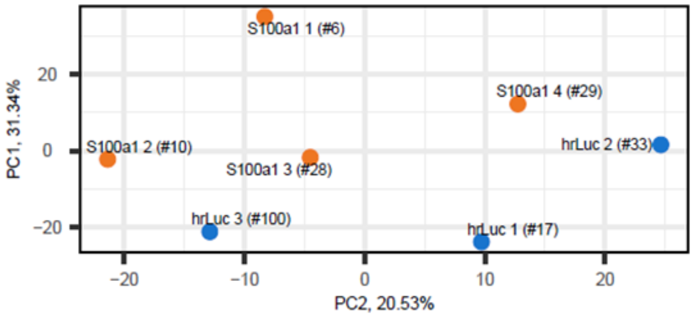

**Appendix figure D** shows separation of AAV5-*hs100A1 (S100a1)* and AAV5-*luc* treated post-MI animal RNAseq. data sets by principal component (PC1 and PC2) visualization.

**Appendix figure E**

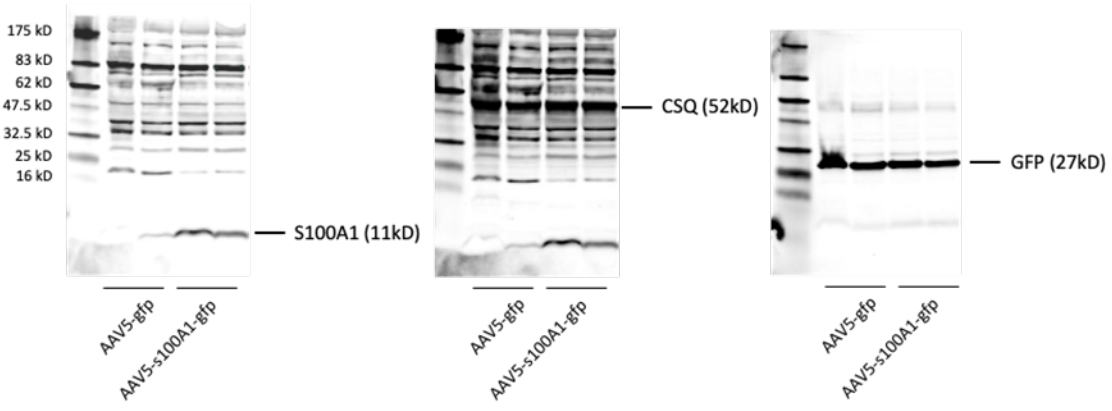

**Appendix figure E** depicts representative immunoblots for anti-S100A1, anti-calsequestrin (CSQ) and anti-GFP protein of LV homogenates of two rAAV5-*gfp* and rAAV5-*hS100A1-gfp* treated post-MI mice. rAAV5-*hS100A1-gfp* treatment resulted in robust S100A1 protein overexpression compared to rAAV5-*gfp* animals after 4 weeks (left). Treatment with intramyocardial injections of $2\times10^{11}$ vgcs in each group yielded comparable GFP protein expression in both groups (middle). CSQ protein staining ensured similar loading of samples (right). The same membrane was used for sequential staining employing appropiate secondary antibodies for near-infrared fluorescent imaging.

### References

1. Weber C, Neacsu I, Krautz B et al. Therapeutic safety of high myocardial expression levels of the molecular inotrope S100A1 in a preclinical heart failure model. *Gene Ther.* 2014;21:131-8.
2. Pleger ST, Shan C, Ksienzyk J et al. Cardiac AAV9-S100A1 gene therapy rescues post-ischemic heart failure in a preclinical large animal model. *Sci Transl Med.* 2011;3:92ra64.
3. Raake PW, Schlegel P, Ksienzyk J et al. AAV6.betaARKct cardiac gene therapy ameliorates cardiac function and normalizes the catecholaminergic axis in a clinically relevant large animal heart failure model. *Eur Heart J.* 2013;34:1437-47.
4. Most P, Seifert H, Gao E et al. Cardiac S100A1 protein levels determine contractile performance and propensity toward heart failure after myocardial infarction. *Circulation.* 2006;114:1258-68.
5. aus dem Siepen F, Buss SJ, Messroghli D et al. T1 mapping in dilated cardiomyopathy with cardiac magnetic resonance: quantification of diffuse myocardial fibrosis and comparison with endomyocardial biopsy. *Eur Heart J Cardiovasc Imaging.* 2015;16:210-6.
6. Riffel JH, Schmucker K, Andre F et al. Cardiovascular magnetic resonance of cardiac morphology and function: impact of different strategies of contour drawing and indexing. *Clin Res Cardiol.* 2019;108:411-429.
7. Raake PWJ, Barthelmes J, Krautz B et al. Comprehensive cardiac phenotyping in large animals: comparison of pressure-volume analysis and cardiac magnetic resonance imaging in pig post-myocardial infarction systolic heart failure. *Int J Cardiovasc Imaging.* 2019;35:1691-1699.
8. Muller T, Boileau E, Talyan S et al. Updated and enhanced pig cardiac transcriptome based on long-read RNA sequencing and proteomics. *J Mol Cell Cardiol.* 2021;150:23-31.
9. Jakobi T, Siede D, Eschenbach J et al. Deep Characterization of Circular RNAs from Human Cardiovascular Cell Models and Cardiac Tissue. *Cells.* 2020;9.
10. Langfelder P, Horvath S. WGCNA: an R package for weighted correlation network analysis. *BMC Bioinformatics.* 2008;9:559.
11. Team RC. R: A language and environment for statistical computing. R Foundation for Statistical Computing, Vienna, Austria., 2022.
12. https://cran.r-project.org/web/packages/fastqcr/index.html.

13. FastQC: A Quality Control Tool for High Throughput Sequence Data

14. Liao Y, Smyth GK, Shi W. The R package Rsubread is easier, faster, cheaper and better for alignment and quantification of RNA sequencing reads. *Nucleic Acids Res.* 2019;47:e47.

15. M C. org.Ss.eg.db: Genome wide annotation for Pig. R package version 3.8.2., 2019.

16. M C. org.Hs.eg.db: Genome wide annotation for Human. R package version 3.8.2. . 2019.

17. BLAST® Command Line Applications User Manual [Internet]. Bethesda (MD): National Center for Biotechnology Information (US); 2008-. Available from: https://www.ncbi.nlm.nih.gov/books/NBK279690/.

18. Hahsler M, Nagar A. rBLAST: R Interface for the Basic Local Alignment Search Tool. R package version 0.99.2., 2019.

19. Love MI, Huber W, Anders S. Moderated estimation of fold change and dispersion for RNA-seq data with DESeq2. *Genome Biology.* 2014;15:550.

20. Zhang B, Horvath S. A general framework for weighted gene co-expression network analysis. *Stat Appl Genet Mol Biol.* 2005;4:Article17.

21. Marini F, Binder H. pcaExplorer: an R/Bioconductor package for interacting with RNA-seq principal components. *BMC Bioinformatics.* 2019;20:331.

22. Ludt A, Ustjanzew A, Binder H, Strauch K, Marini F. Interactive and Reproducible Workflows for Exploring and Modeling RNA-seq Data with pcaExplorer, Ideal, and GeneTonic. *Curr Protoc.* 2022;2:e411.

23. Blighe K LA. PCAtools: PCAtools: Everything Principal Components Analysis. R package version 2.10.0, https://github.com/kevinblighe/PCAtools. 2022.

24. Wu T, Hu E, Xu S et al. clusterProfiler 4.0: A universal enrichment tool for interpreting omics data. *The Innovation.* 2021;2:100141.

25. Yu G, Wang LG, Han Y, He QY. clusterProfiler: an R package for comparing biological themes among gene clusters. *Omics.* 2012;16:284-7.

26. Yu G, He QY. ReactomePA: an R/Bioconductor package for reactome pathway analysis and visualization. *Mol Biosyst.* 2016;12:477-9.

27. Langfelder P, Horvath S. WGCNA: an R package for weighted correlation network analysis. *BMC Bioinformatics.* 2008;9:559.

28. Langfelder P, Zhang B, Horvath S. Defining clusters from a hierarchical cluster tree: the Dynamic Tree Cut package for R. *Bioinformatics.* 2007;24:719-720.

29. Gao E, Lei YH, Shang X et al. A novel and efficient model of coronary artery ligation and myocardial infarction in the mouse. *Circ Res.* 2010;107:1445-53.

30. Takagawa J, Zhang Y, Wong ML et al. Myocardial infarct size measurement in the mouse chronic infarction model: comparison of area- and length-based approaches. *J Appl Physiol (1985).* 2007;102:2104-11.
